# Exploring PBP2a resistance in MRSA by comparison between molecular covalent docking and non-covalent docking

**DOI:** 10.1101/2025.02.20.639213

**Authors:** Ye-Qiang Cao, Yi-Feng Shi

## Abstract

**Background and objectives:** The presence of penicillin-binding protein 2a (PBP2a) is the cause of Methicillin-resistant *Staphylococcus aureus* (MRSA), which is important nosocomial pathogens worldwide and now are also of growing importance in community-acquired infection. The PBP2a resistance depends upon a supplementary peptidoglycan transpeptidase, which continues to function when normal PBPs have been inactivated by beta-lactam antibiotics. Analysis and quantitative study of the molecular interactions of PBP2a against β-lactam antibiotics are required, as they support explaining and enhance understanding of the structure-activity relationship of antibiotic resistance.

**Methods:** Bioinformatics and computational methods have been highly effective tools for β-lactams targeting PBPs to tackle the urgent threat of antimicrobial resistance. Regarding β-lactam antibiotics targeting PBP2a and PBPs, we applied different docking programs to illustrate inhibition mode, MM/GBSA to estimate the binding free energies, and molecular dynamic simulation to validate and analyze the molecular interactions.

**Results:** Based on β-lactam antibiotics targeting PBPs as covalent inhibitors, covalent docking was employed to provide explicit models of PBP2a against susceptible β-lactam antibiotics. The simultaneous use of non-covalent docking enhances our comprehensive comprehension of the resistance of PBP 2a, which resulted from the lack of covalent linked to β-lactam antibiotics. The selected antibiotics strongly interact with PBP2a, revealing the essential amino acid residues and binding affinity for inhibition. MD simulations were performed for the ligand-bound state of PBP-2a to explain their interaction and conformational changes. These findings are also strongly supported by root-mean-square deviation (RMSD), root-mean-square fluctuation (RMSF) and Hydrogen bond analysis of the protein-ligand complex.

**Conclusions:** Our research offers extensive knowledge of the PBP2a-lactam interactions for the ability of known antibiotics to combat MRSA. The simulation results indicating stability and accuracy provide valuable insights for the advancement of pharmaceutical interventions against infectious diseases

Figure abstract:
The interaction of Methicillin with PBP2a of MRSA.
The protein is shown as a cartoon model, and the covalent binding of the ligand and serine active site is shown as a stick model.

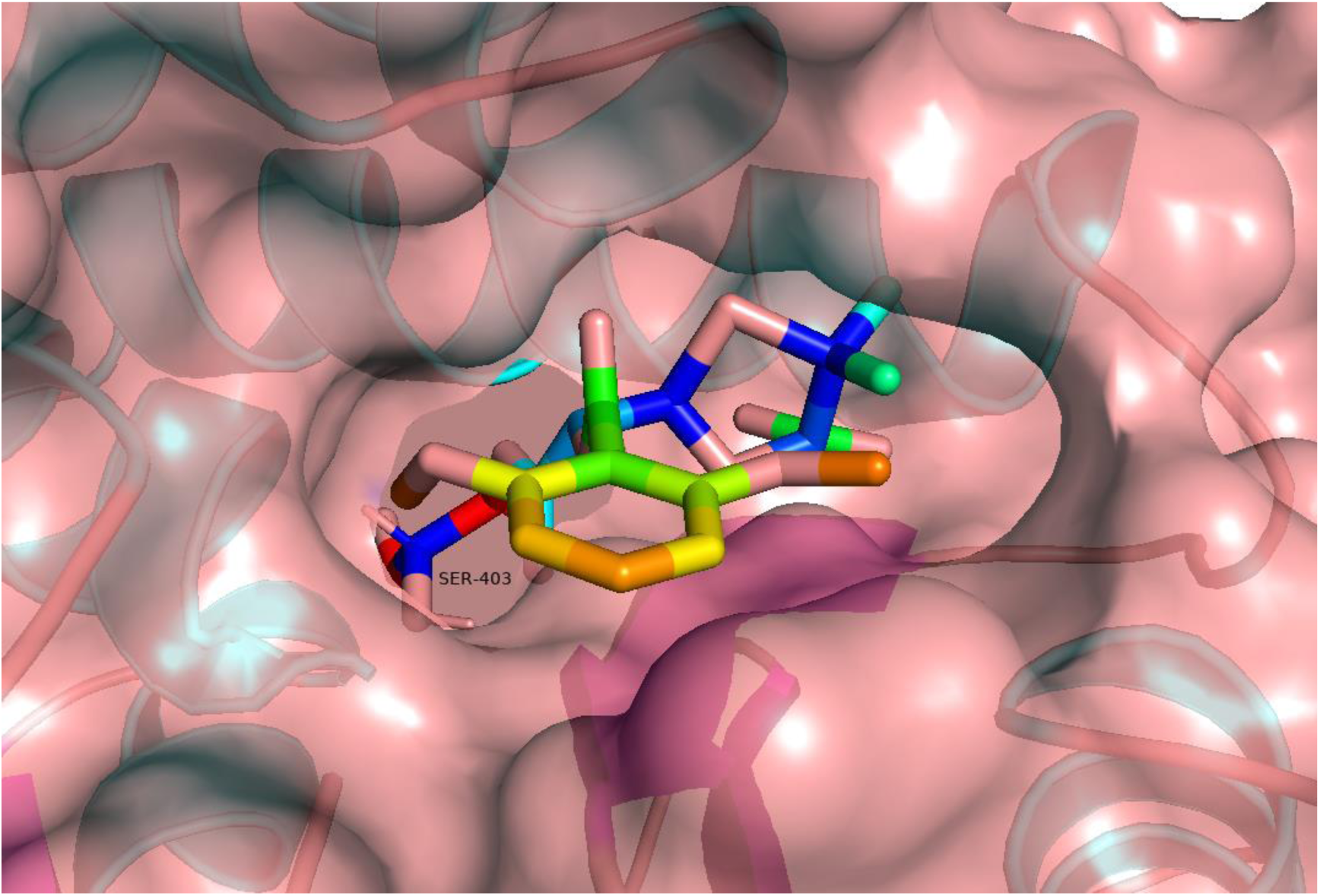

## 1. Introduction

*Staphylococcus aureus* has been an exceptionally successful pathogen owing to its abundance (in the environment and the normal flora) and the variety of virulence factors that it possesses. *S. aureus* may colonize various mucosal sites of the body and is an important causative agent of invasive infections in modern age medicine. Methicillin-resistant *S. aureus* (MRSA) strains—first described in 1961—are characterized by an altered penicillin-binding protein (PBP2a) and resistance to all penicillins, cephalosporins, and carbapenems, which may add significant roles in other pathologies (epidural abscesses, meningitis, toxic shock syndrome (TSS), urinary tract infections (UTIs), septic thrombophlebitis, etc.). MRSA initially is largely a nosocomial infection, leading to so-called hospital-associated MRSA (hospital-associated MRSA, HA-MRSA). The epidemiology of MRSA infections shifted during the 1990s when community-associated MRSA (CA-MRSA) emerged. Additionally, MRSA can be harbored by livestock (livestock-associated MRSA, LA-MRSA), first described at the beginning of the 2000s, where it can cause disease in those animals and also spread to humans via contact. According to the official assessments (CDC) based on various factors, such as treatability, mortality, burden on the healthcare infrastructure and the community, prevalence and increasing trends of resistance, preventability and transmissibility, in addition to the drugs that are currently in the pipeline, MRSA has been classified as a serious threat. ^[1∼4]^

Penicillin-binding proteins (PBPs) are a family of membrane-bound proteins that are essential for the synthesis of peptidoglycan, the main component of the cell wall. PBPs are serine acyltransferases that facilitate the production of cross-linked peptidoglycan and serve as targets for β-lactam antibiotics due to their transpeptidase-related catalytic activity. β-lactam antibiotics covalently bind to the active site serine of PBPs after adhering to the PBP catalytic cleavage, resulting in the formation of a slowly hydrolyzed acyl-enzyme complex that reduces peptidoglycan cross-linking, leading to cell lysis. *S. aureus* normally possesses four endogenous PBPs, PBP1, PBP2, PBP3, and PBP4, with an additional, acquired PBP2a found in MRSA. PBP2a is considered a surrogate enzyme, taking over the cell wall synthesis task from the normal complement of *S. aureus* PBPs. It has a poor affinity for β-lactam antibiotics and exhibits transpeptidase activity, crucial for peptidoglycan biosynthesis. PBP2 is the primary target for β-lactam antibiotics, and PBP2a confers resistance to all currently available β-lactam medications. When interacting with β-lactam antibiotics, these enzymes undergo significant conformational shifts that may affect PBP2a’s catalytic activities in bacterial cell wall cross-linking. The diversity of PBPs and their varying sensitivities to β-lactams add to the complexity of the mechanism of bacterial antibiotic resistance. ^[5∼8]^

The use of computational methods in drug design has increased rapidly in recent years, and numerous tools are available to support this process. Virtual screening is a cost-effective and time-saving method that helps researchers narrow down the number of compounds for further experimental analysis.^[9∼11]^ To identify promising lead compounds, large libraries of commercially available small molecules perform docking-based virtual screening. Verma et al.^[12]^ and Alhadrami et al.^[13]^ tested an in-house library of common plant-based phenolic compounds and found some flavonoid compounds effective inhibiting Penicillin Binding Protein-2a (PBP-2a) in MRSA using molecular docking. Farid et al. reproposed the antibacterial potential of Amphotericin B, an anti-fungal drug, against MRSA revealed by antimicrobial screening, molecular docking, and mode of action analysis targeting Penicillin Binding Protein 2a^[14]^. Qureshi et al. designed new quinazolinone analogs effective against MRSA strains, guided by molecular docking, In-silico and MM-GBSA study^[15]^.

Molecular docking allows the underlying mechanism of inhibition and the binding pose adopted by the compound in the active site of its target. Molecular dynamics (MD) simulations provide detailed insights into the physical movements and interactions of atoms and molecules in a system.^[16.17]^ Tripathi et al.^[18]^ verified the molecular interaction of Penicillin binding protein-2 *Neisseria meningitidis,* the main causative agents of bacterial meningitis, to different generations of β-lactam antibiotics and concluded that the third generation of β-lactam antibiotics shows efficient binding with Penicillin binding protein-2 of *N. meningitidis*. Kumar et al.^[19]^ performed molecular docking studies to explore possible binding modes of Penicillin derivatives and Cephalosporins into all types of PBPs for upper respiratory tract bacterial pathogens and revealed that the Cephalosporins show higher affinity with PBPs than the Penicillin derivatives, especially the fifth-generation Cephalosporins, Ceftobiprole and Ceftaroline. Alshabrmia et al.^[20]^ studied the mechanisms of binding interactions between wild-type and mutated PBP2a against Cefoxitin using molecular docking and MD simulations. Their result reveals that the binding of mutated PBP2a shows resistance and unstable behavior by binding with Cefoxitin.

Covalent drugs such as penicillin and aspirin have been used to treat diseases for more than a century, but their mechanisms of covalent action were unknown until the discovery of containing protein-reactive functional groups. The continuous approval of covalent drugs in recent years for the treatment of diseases has led to an increased search for covalent agents by medicinal chemists and computational scientists worldwide^[21, 22]^. Most computational programs for covalent docking rely on directly linking models to model conformations under the constraint of a predefined bond between a ligand and a corresponding amino acid site. CovDock is based on the Schrӧdinger Glide docking algorithm and Prime structure refinement methodology^[23,24]^. CovDock uses traditional non-covalent docking approaches to dock a ligand to a protein target, and then models the covalent bond attachment and refines the complex ^[25, 26]^. In this study, comparing covalent docking and non-covalent docking, validated by Molecular dynamics simulation, provided useful information on binding modes of a series of known β−lactam antibiotics for investigating PBP2a resistance mechanism in MRSA. We sought to demonstrate covalent docking and the existence of suitable programs for the rational docking pursuit when dealing with covalent targets.

## 2. Materials and Methods

### 2.1 Selection of protein and sequence alignment

Three PBPSs, PBP2A, PBP2 and PBP3 were selected for this study. The three-dimensional structures (3D) and sequences of the PBPs were obtained from the Protein Structure Data Bank (PDB), and the penicillin-binding protein 2a (PDB ID: 1MWU) of *Staphylococcus aureus* in MRSA^[27]^, as well as the penicillin-binding protein 2 (PDB ID: 7KIS) of *Pseudomonas aeruginosa* WCK 5153^[28]^ and the penicillin-binding protein 3 (PDB ID: 3PBR) of *Pseudomonas aeruginosa* PAO 1^[29]^ were searched in the RCSB PDB database. The multiple sequence alignment of PBPs was performed using MEGA11.0 software^[10]^.

### 2.2 Protein receptor preparation

Co-crystalline ligands were identified and removed from the target protein, followed by removal of crystal water molecules from the 3D coordinate file to avoid solvent-mediated salt-bridge interactions of ligands with the active and allosteric sites of the receptor. The Build module of Schrodinger Maestro 12.8 was used to add hydrogen and reconstruct the missing side chains into the target Protein structure. Protein Preparation Wizard was used for protonation and energy minimization, and the pH of amino acid residue matrix was set to 7.0. The energy minimization position function of OPLS4 force field algorithm was used to minimize the energy of the receptor and refine the 3D structure.

### 2.3 Ligand acquisition and preparation

The structures of β-lactam antibiotics were obtained from NCBI PubChem Compound database, and the structures were drawn using Chemsketch, hydrogen atoms were added to all structures, and atomic charges were calculated, using the Ligprep module of Schrodinger Maestro 12.8, The pH of the ionized state was set to 7.0, and the molecules were energy minimized using the OPLS3 force field. During docking, the flexible conformation of the compound is geometrically optimized using the chimera conformation. In this study, the general name, molecular weight, chemical formula and two-dimensional structure list of β-lactam antibiotics, as shown in Table 1, were used to screen 13 known potential antibiotics for molecular docking study of the functional residues of penicillin-binding protein.

**Table 1.**
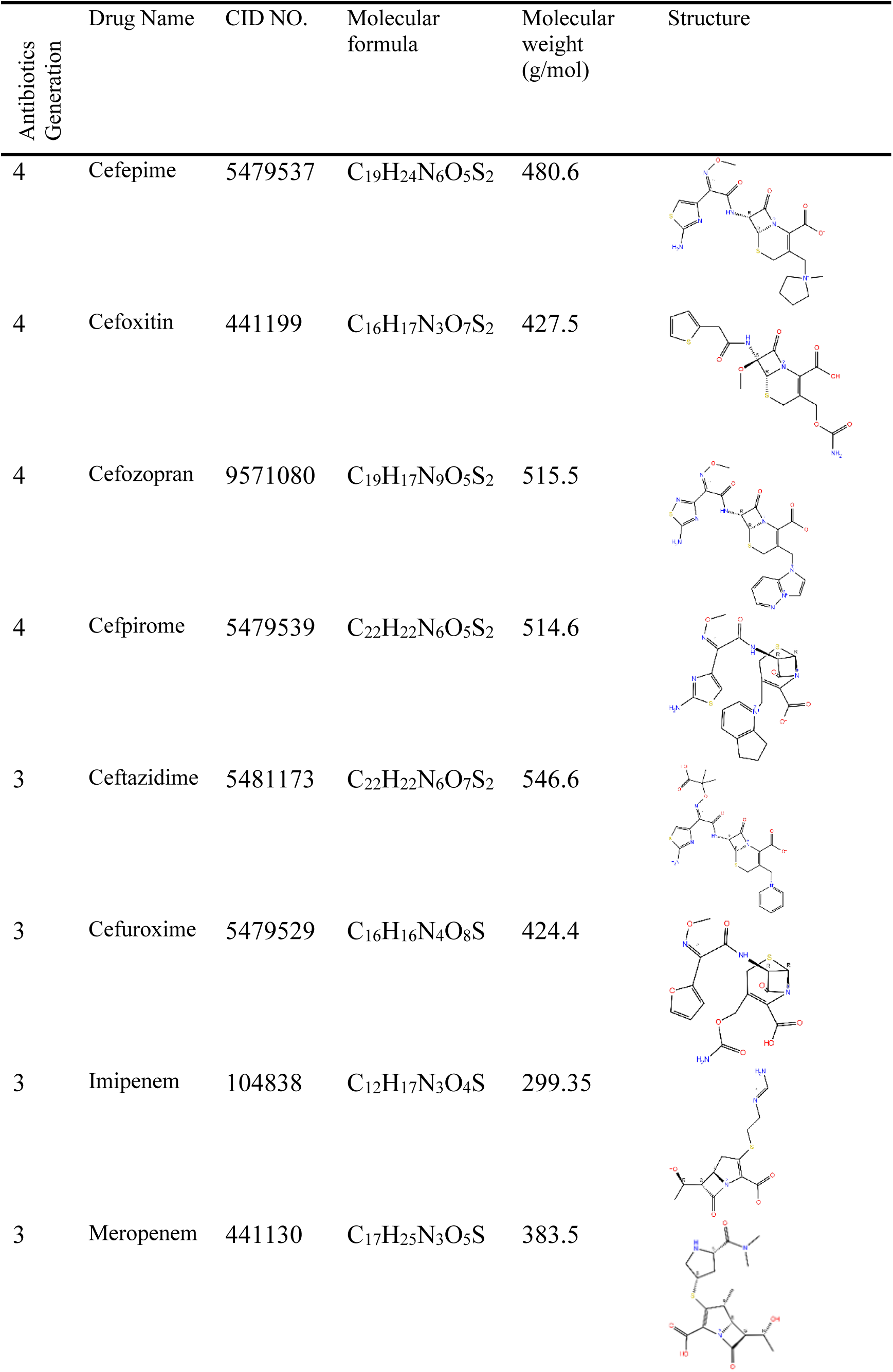
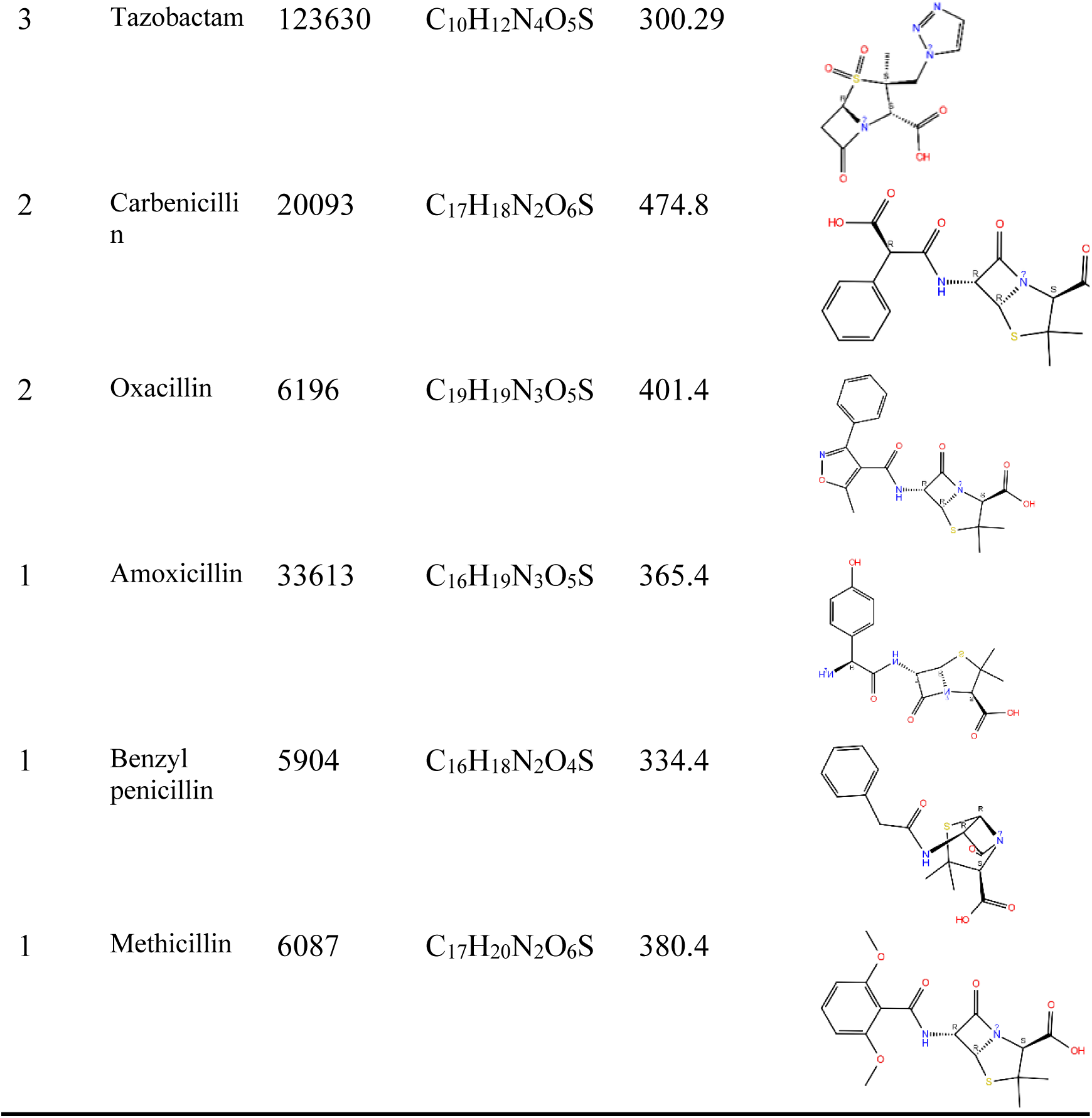
β-Lactam antibiotics used in this study for docking ligands.

| Antibiotics<br>Generation | Drug Name | CID NO. | Molecular<br>formula | Molecular<br>weight<br>(g/mol) | Structure |
| --- | --- | --- | --- | --- | --- |
| 4 | Cefepime | 5479537 | C<sub>19</sub>H<sub>24</sub>N<sub>6</sub>O<sub>5</sub>S<sub>2</sub> | 480.6 |  |
| 4 | Cefoxitin | 441199 | C<sub>16</sub>H<sub>17</sub>N<sub>3</sub>O<sub>7</sub>S<sub>2</sub> | 427.5 |  |
| 4 | Cefozopran | 9571080 | C<sub>19</sub>H<sub>17</sub>N<sub>9</sub>O<sub>5</sub>S<sub>2</sub> | 515.5 |  |
| 4 | Cefpirome | 5479539 | C<sub>22</sub>H<sub>22</sub>N<sub>6</sub>O<sub>5</sub>S<sub>2</sub> | 514.6 |  |
| 3 | Ceftazidime | 5481173 | C<sub>22</sub>H<sub>22</sub>N<sub>6</sub>O<sub>7</sub>S<sub>2</sub> | 546.6 |  |
| 3 | Cefuroxime | 5479529 | C<sub>16</sub>H<sub>16</sub>N<sub>4</sub>O<sub>8</sub>S | 424.4 |  |
| 3 | Imipenem | 104838 | C<sub>12</sub>H<sub>17</sub>N<sub>3</sub>O<sub>4</sub>S | 299.35 |  |
| 3 | Meropenem | 441130 | C<sub>17</sub>H<sub>25</sub>N<sub>3</sub>O<sub>5</sub>S | 383.5 |  |
| 3 | Tazobactam | 123630 | $C_{10}H_{12}N_4O_5S$ | 300.29 | |
| 2 | Carbenicillin | 20093 | $C_{17}H_{18}N_2O_6S$ | 474.8 | |
| 2 | Oxacillin | 6196 | $C_{19}H_{19}N_3O_5S$ | 401.4 | |
| 1 | Amoxicillin | 33613 | $C_{16}H_{19}N_3O_5S$ | 365.4 | |
| 1 | Benzyl penicillin | 5904 | $C_{16}H_{18}N_2O_4S$ | 334.4 | |
| 1 | Methicillin | 6087 | $C_{17}H_{20}N_2O_6S$ | 380.4 | |

### 2.4 Non-covalent approach of molecular docking

Glide module (Schrödinger, LLC, NY) was used to perform the docking studies^[30, 31]^. The docking study comprises five steps ligand preparation, protein preparation, receptor grid generation, actual docking procedure and finally viewing the docking results using the pose-viewer.

The active site was defined by removing the cocrystallized ligand and generating the grid box at the centroid of the workspace ligand as selected in the receptor folder. Ligand Docking OPLS 2005 force field was used during docking. The receptor grid generated previously was selected for the docking of ligands prepared using Lig prep. Standard Precision (SP) feature was used to perform flexible docking. The van der Waals radii were scaled to 0.80 default and default partial cutoff charge of 0.15 to decrease the penalties for close contacts. No constraints were set to define ligand-receptor interactions. The structure output form was set to pose viewer file so as to view the output of the resulting docking studies from the pose-viewer.

Pose-Viewer was used to see docking results. The H-bonding and salt bridge interaction between ligands and receptors were visualized using default settings to analyze the binding modes of the ligands to the receptor. No constraints were set to define ligand-receptor interactions. A computer model score system that encompasses the grid score, proprietary GLIDE score, and the internal energy strain were used to rank the final ligand binding poses. The score function of Glide, or Glide score, was used for binding affinity prediction and ligand ranking.

### 2.5 Covalent approach of molecular docking

Ligands were covalently docked against different PBPs via a β-lactam ring-opening reaction using Schrodinger’s CovDock^[32, 33]^. CovDock enables covalent docking of small molecules to proteins by first mutating the reactive residue to alanine, docking the ligand to the active site using extensive conformational sampling with certain translational constraints (to prevent the reactive payload from moving away from the binding residue), and then mutating the reactive residue back to its original state and building the covalent bond to the ligand’s payload. The complex-bound ligand is then minimized and binding energies are predicted using a scoring function that takes into account both the affinity of the initial docking and the conformational strain associated with the covalent binding and subsequent active site rearrangements (ligands that require significant structural shifts to accommodate receive less favorable scores). Serine 403 was defined as the reactive residue in PBP2a, based upon significant evidence from the literature. The thoroughness was set to “pose prediction” for increased accuracy.

### 2.7 Binding free energy calculation using Prime/MM-GBSA approach

The approach of molecular mechanics, along with generalized Born and surface area solvation (MM-GBSA)^[34, 35]^, is applied to estimate the absolute ligand–protein binding affinities by calculating its free binding energy. The prime module containing the MM-GBSA panel with default settings was used for estimating the free-binding energy. The docked molecules-receptor complex with a top Glide XP docking score was considered to perform this MM/GBSA task. The formula employed for computing the binding free energies of the ligands is mentioned below.

ΔG_(binding)_ = ΔE_(MM)_ +ΔG_(sovaltion)_ +ΔG_(SA)_

MM/GBSA_(ΔGbind)_ = G_complex(minimized)_ - [G_ligand (unbound,minimized)_ + G_receptor (unbound, minimized)_ ]

where ΔG_(binding)_ is mainly the difference between the polar desolvation energies, ΔE_(MM)_ is basically the difference between the minimized energies of the PBP docked complex, and the sum of the individually calculated minimized energies of the PBP protein structure and the corresponding hit. Similarly, ΔG_(solvation)_ is the difference in the GBSA solvation energy of the ligand-PBP complex and the summation of the solvation energies of the PBP structure and ligand. Whereas ΔG_(SA)_ is a change in the surface area energies of the bounded complex and the total surface area energies of the unaligned receptor and picked hits. The solvation model of VSGB (variable-dielectric generalized Born) combined with the variable-dielectric generalized Born model under the OPLS4 force field calculated the ligand binding energy in the Prime MM-GBSA method.

Where MM/GBSA_ΔGbind_ is the calculated relative free energy that includes both ligand and receptor strain energy. G_complex(minimized)_ is the MM/GBSA energy of the minimized complex, and G_ligand (unbound, minimized)_ is the MM/GBSA energy of the ligand after removing it from the complex and allowing it to relax. G_receptor (unbound, minimized)_ is the MM/GBSA energy of protein after separating it from the ligand.

### 2.8 Molecular dynamics (MD) simulation

The molecular dynamic simulation study was performed to examine the conformational changes in the protein that occurred due to the ligand-binding site and to evaluate the effect of these changes over the protein-ligand complex. The PBPs docking complex underwent a 100 ns molecular dynamics (MD) simulation using Desmond (Schrödinger LLC) software^[36∼38]^. The complex was placed in the protein preparation wizard in Maestro for optimization, analysis, and refinement of the docking complex. Before the simulations, the docking complexes were optimized and minimized by applying Protein Preparation Wizard. All systems were set up via the System Builder tool, utilizing the TIP3P solvent model in an orthorhombic box. The OPLS 2005 force field was chosen in the simulations, and counter ions were introduced to neutralize the models. To replicate physiological conditions, sodium chloride (NaCl) was added to achieve a final concentration of 0.15 M, providing Na ^+^ and Cl − ions. The NPT ensemble with a temperature of 300 K and pressure of 1 atm was chosen for the entire simulation period of 100 nsec. The models were equilibrated before starting the simulation. Trajectories were saved for analysis every 100 ps, and the stability of the simulation was monitored by comparing the root mean square deviation (RMSD) of the protein over time (Maiorov associated with Desmond popular as Simulations Interactions Diagram (SID) was put into practice to scrutinize the RMSD, RMSF and other critical details from the simulated complexes. The root mean square deviation (RMSD) analysis, root mean square fluctuation (RMSF), full contacts, and the interaction fractions maintained throughout the MD simulation indicate the protein’s stability and ligand’s stability in bound form.

## 3. Results and discussion

### 3.1 Docking of methicillin and oxacillin into PBP2a and PBPs

To evaluate PBP2a resistance in MRSA, Penicillin-binding protein 2a from methicillin-resistant *Staphylococcus aureus* strains, penicillin-binding protein 2 from *Pseudomonas aeruginosa*, and penicillin-binding protein 3 from *Pseudomonas aeruginosa* Meropenem complex were selected and their sequence alignment was performed (Fig. 1). In particular, serine (S403) is the active site core, and its hydroxyl group is strongly nucleophilic, setting the stage for subsequent chemical reactions. When β-lactam antibiotics, such as penicillin, are in close proximity to PBPs, the carboxyl group in the β-lactam ring of the antibiotic covalently binds to the serine hydroxyl group to form a serine ester, which irreversibly inhibits the activity of transpeptidase and blocks the normal synthesis of the cell wall, resulting in the inhibition of the bacterial growth.

**Figure 1.**
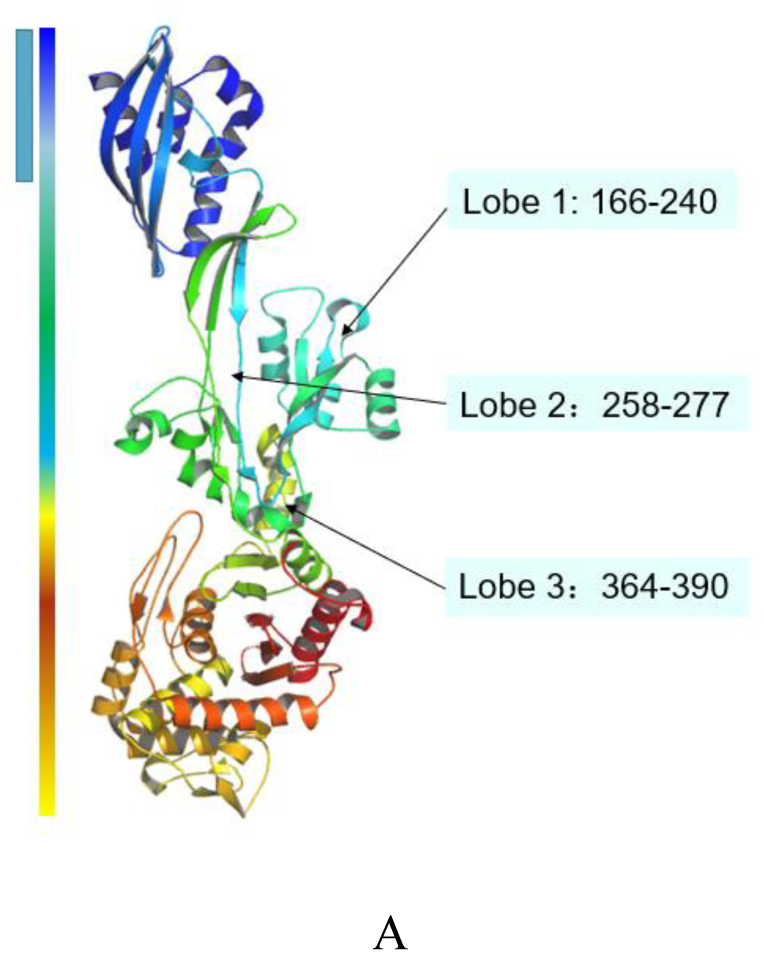
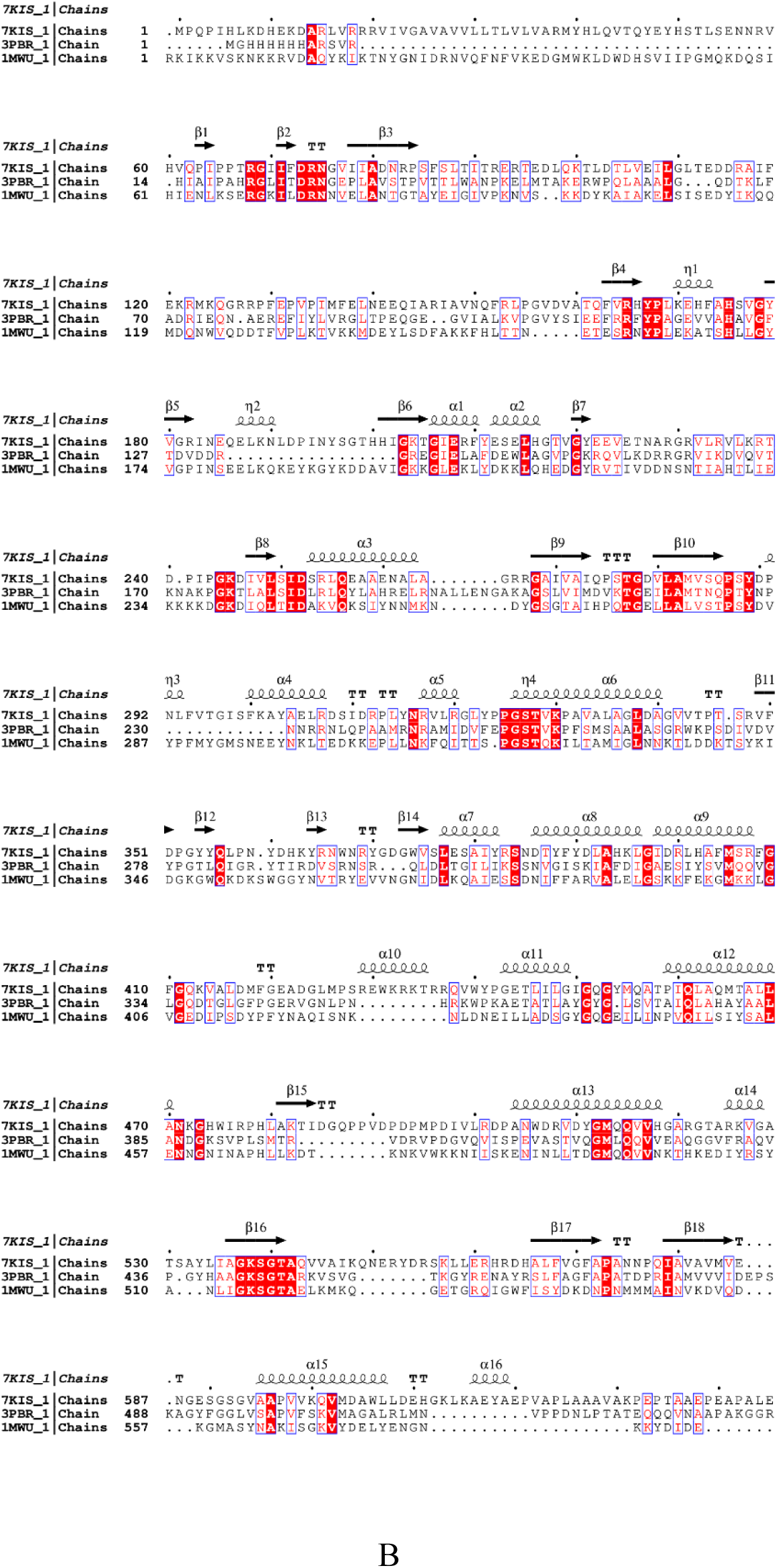
(A)The crystal structure of PBP2a consists of three domains and (B)Sequence alignment between PBP2a(PDB ID: 1MWU), PBP2 (PDB ID: 7KIS) and PBP3(PDB ID: 3PBR).

Covalent docking of methicillin and oxacillin was performed into PBP2a, PBP2 and PBP3 respectively (Figure 2, Table2). For different PBPs, oxacillin had a higher affinity (Table 4), with binding energies of -45.25 kcal/mol, -14.08 kcal/mol, and -71.93 kcal/mol, respectively, which demonstrated that oxacillin inhibited PBPs better than methicillin. Meanwhile, the binding energies of PBP2a were -64.12 kcal/mol and -62.84 kcal/mol when docked with methicillin and oxacillin, respectively, which were lower than those of PBP2 and PBP3, suggesting that its affinity was lower. PBP2a was found to form 5 and 4 hydrogen bonds with methicillin and oxacillin, respectively, PBP2 formed 7 and 5 hydrogen bonds with methicillin and oxacillin, respectively, and PBP3 formed 6 hydrogen bonds with methicillin and oxacillin. PBP2a shows more resistance to methicillin and oxacillin than PBP2 and PBP3.

**Figure 2.**
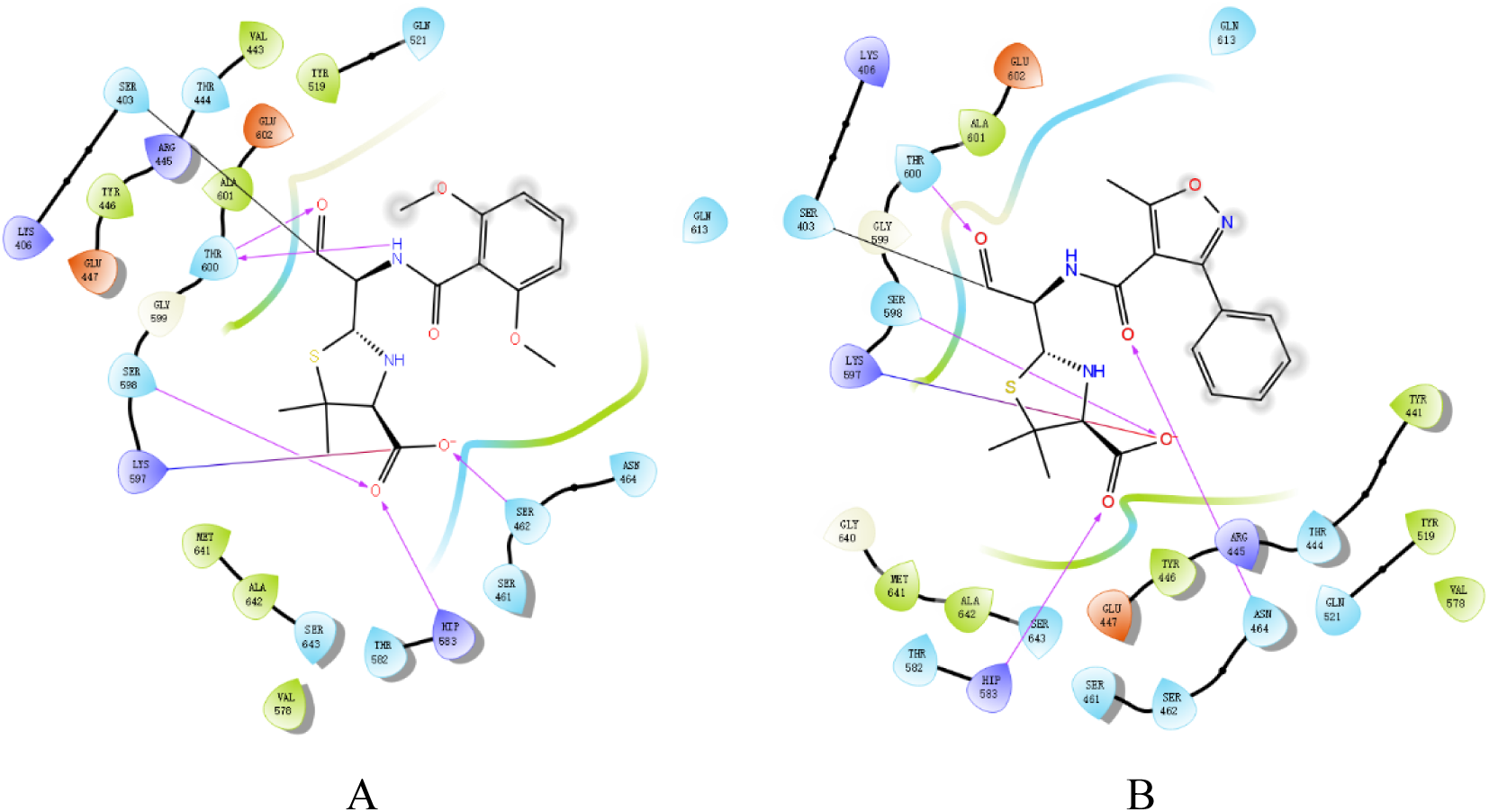
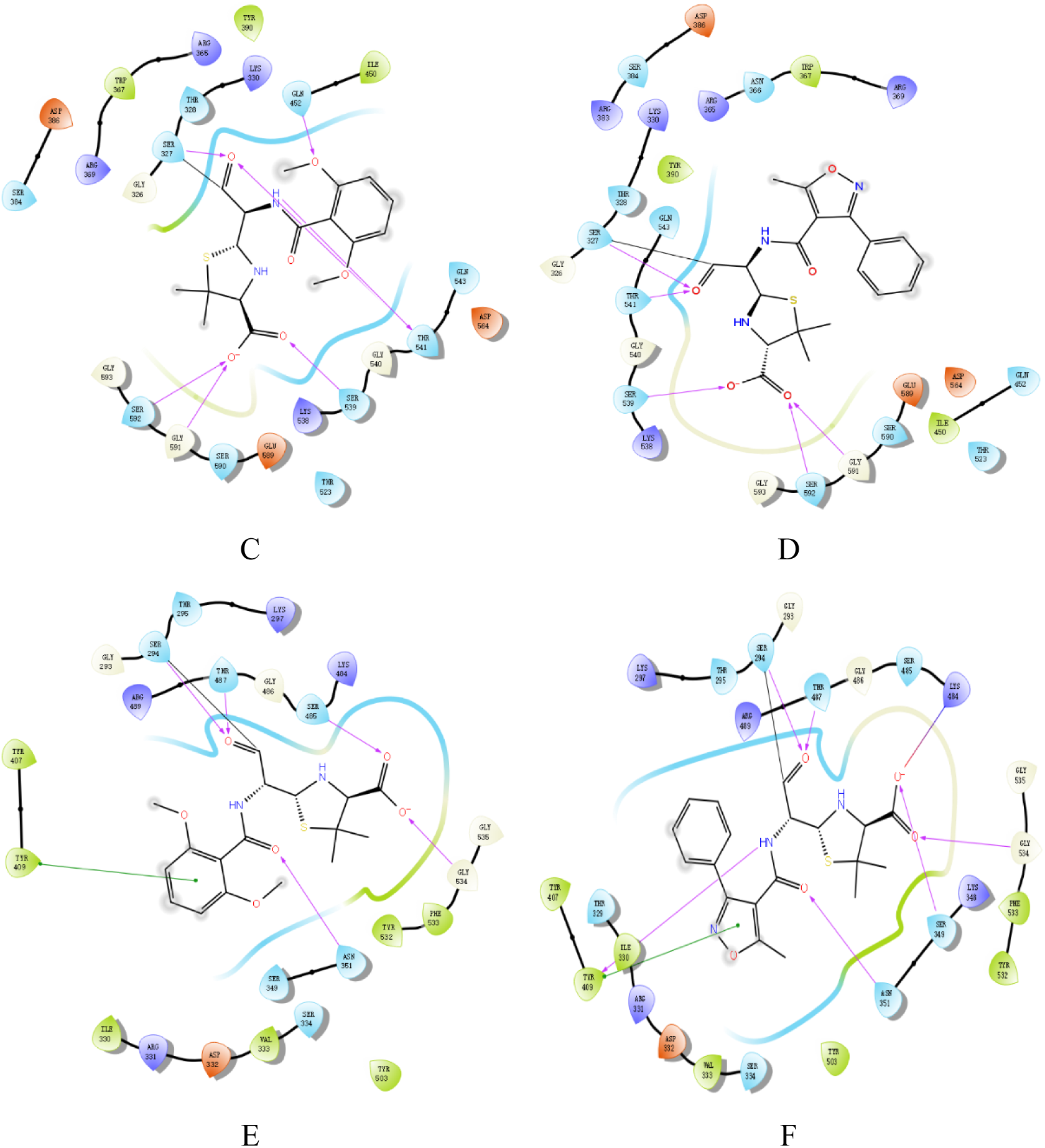
Covalent docking methicillin and oxacillin into PBPs respectively PBP2a(PDB ID: 1MWU), PBP2 (PDB ID: 7KIS) and PBP3(PDB ID: 3PBR). A: Methicillin-7KIS; B: Oxacillin-7KIS; C: Methicillin-1MWU; D: Oxacillin-1MWU; E: Methicillin-3PBR; F: Oxacillin-3PBR

**Table 2.** Interaction patterns of different PBPs with methicillin and oxacillin.

| Ligand–enzyme | Interacting residues | H-bonds | MMGBSA |  |
| --- | --- | --- | --- | --- |
|  |  |  | dG Bind | docking |
|  |  |  | Energy | score |
|  |  |  | (Kcal/mol) |  |
| Methicillin-7KIS | GLN452、THR541(2)、SER539、GLY591、<br>SER592、SER327 | 7 | -34.53 | -5.737 |
| Oxacillin-7KIS | SER327、THR541、SER539、SER592、GLY591 | 5 | -45.25 | -3.988 |
| Methicillin-1MWU | THR600(2)、SER598、HIP583、SER462 | 5 | -64.12 | -5.899 |
| Oxacillin-1MWU | THR600、SER598、HIP583、SER462、 | 4 | -62.84 | -6.313 |
| Methicillin-3PBR | SER294、THR487、SER485、GLY534、ASN351、<br>TYR409、 | 6 | -65.49 | -8.373 |
| Oxacillin-3PBR | SER294、THR487、GLY534、SER349、ASN351、<br>TYR409 | 6 | -71.93 | -2.647 |

### 3.2 Docking of four generations of β-lactam antibiotics into PBP2a

According to β-lactam antibiotics as a suicide inhibitor, we have assumed the mechanism of penicillin-binding proteins (PBPs) resistance that the susceptible activities of β-lactam antibiotics are attributed to their covalent binding affinity to PBPs, while non-covalent binding affinity to PBPs is accountable for their resistance capabilities of β -lactam antibiotics. Therefore, both covalent and non-covalent docking approaches were used to investigate the molecular interactions of PBP2a resistance. By exploring molecular interactions and binding affinities, we aim to uncover promising inhibitors that can effectively hinder the activity of penicillin-binding protein 2a.

Covalent molecular docking was performed on four generations of β-lactam antibiotics with PBP2a using Schringer’s CovDcok. The interaction of these antibiotics with modeled protein was selected based on intermolecular binding and Hydrogen bonding interaction as illustrated in Figure 3. These values along with the hydrogen bond forming residues are presented in Table 2. Binding energies including docking and MM/GBSA for each docked complex are shown in Table 3. The results of non-covalent docking of protein-ligand interactions (Figure 4 and Table 4) and binding energies (Table 5) were obtained by using Schrodinger’s GLIDE docking and Prime/MM-GBSA. Comparison between molecular covalent docking and non-covalent docking into PBP2a is shown in Table 6.

**Figure 3.**
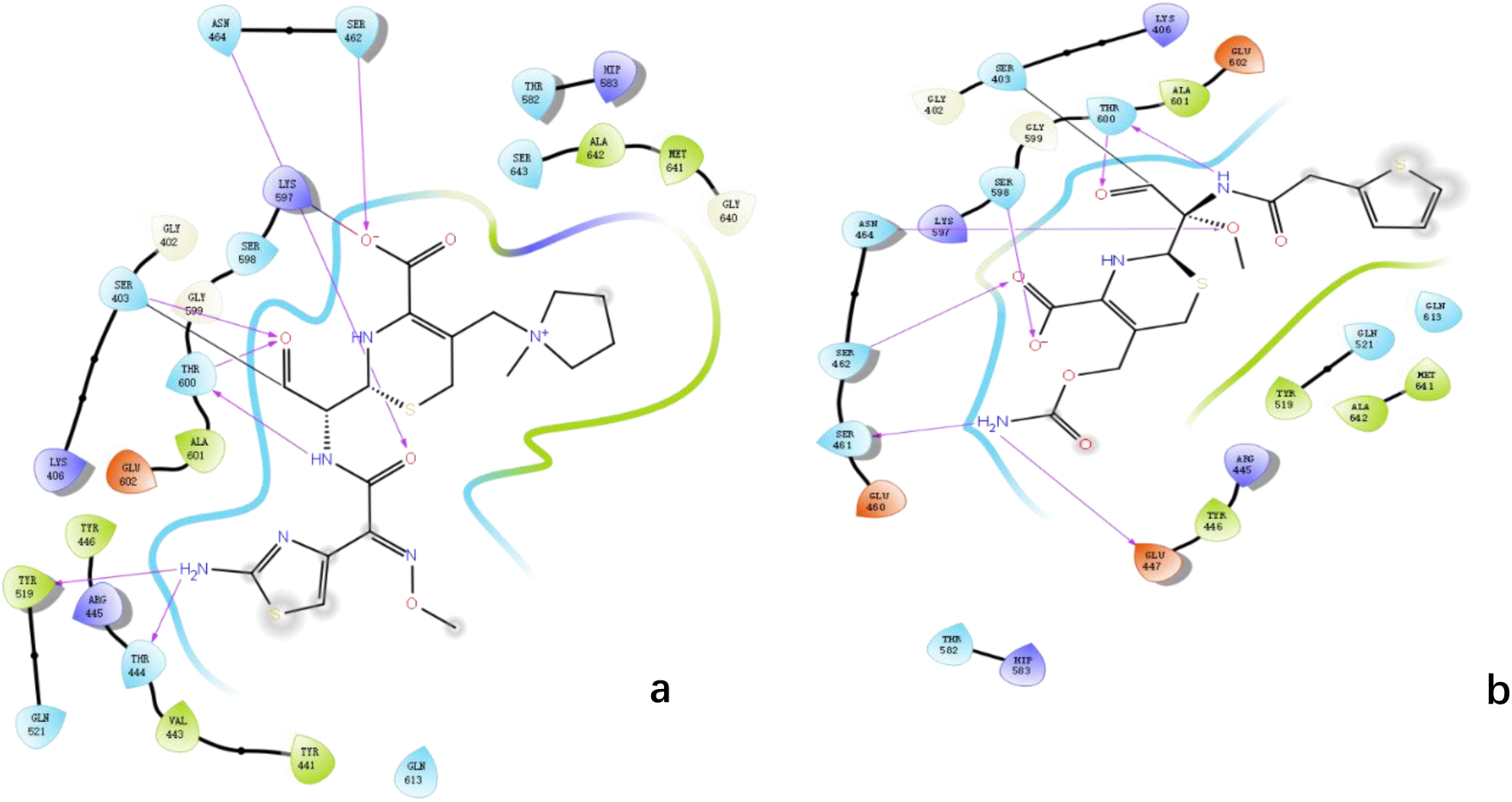
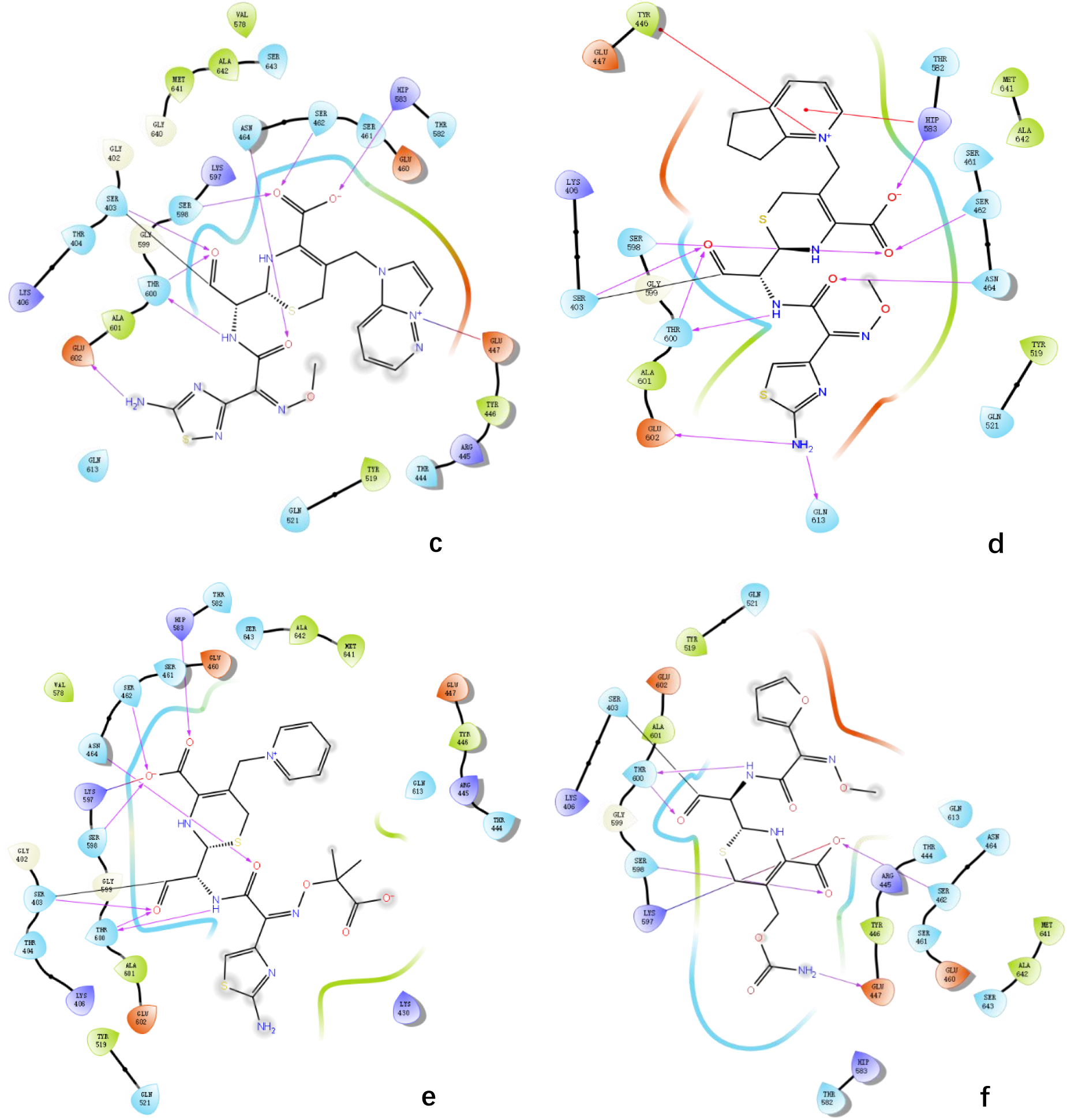
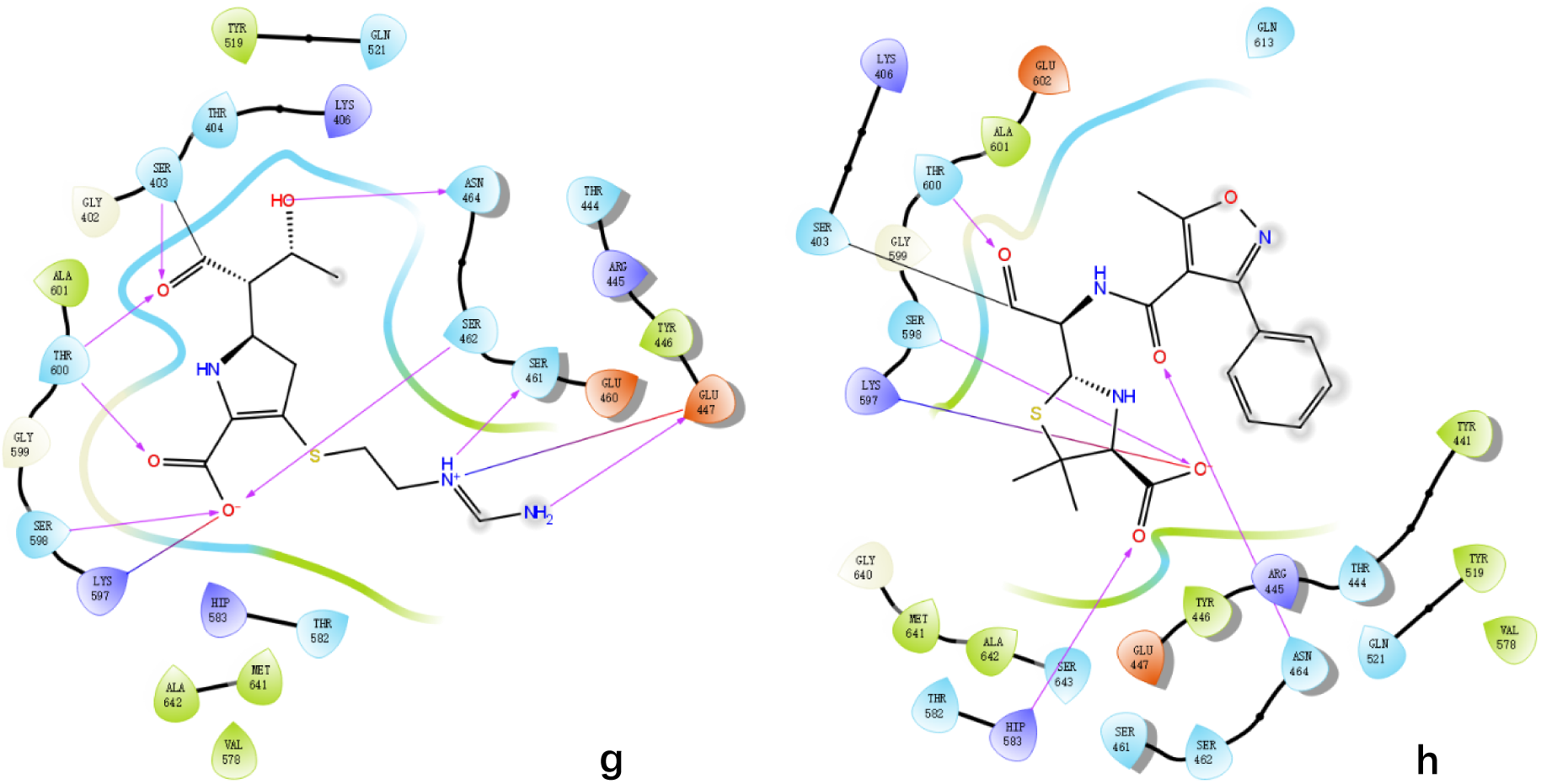
Covalent docking interaction between different generations of β-lactam antibiotics and PBP2a. a |Interaction between 1MWU and Cefepime. b |Interaction between 1MWU and cefoxitin. c |Interaction between 1MWU and Cefozopran. d |Interaction between 1MWU and Cefpirome. e |Interaction between 1MWU and Ceftazidime. f |Interaction between 1MWU and Cefuroxime. g |Interaction between 1MWU and Imipenem. h |Interaction between 1MWU and oxacillin .

**Figure 4.**
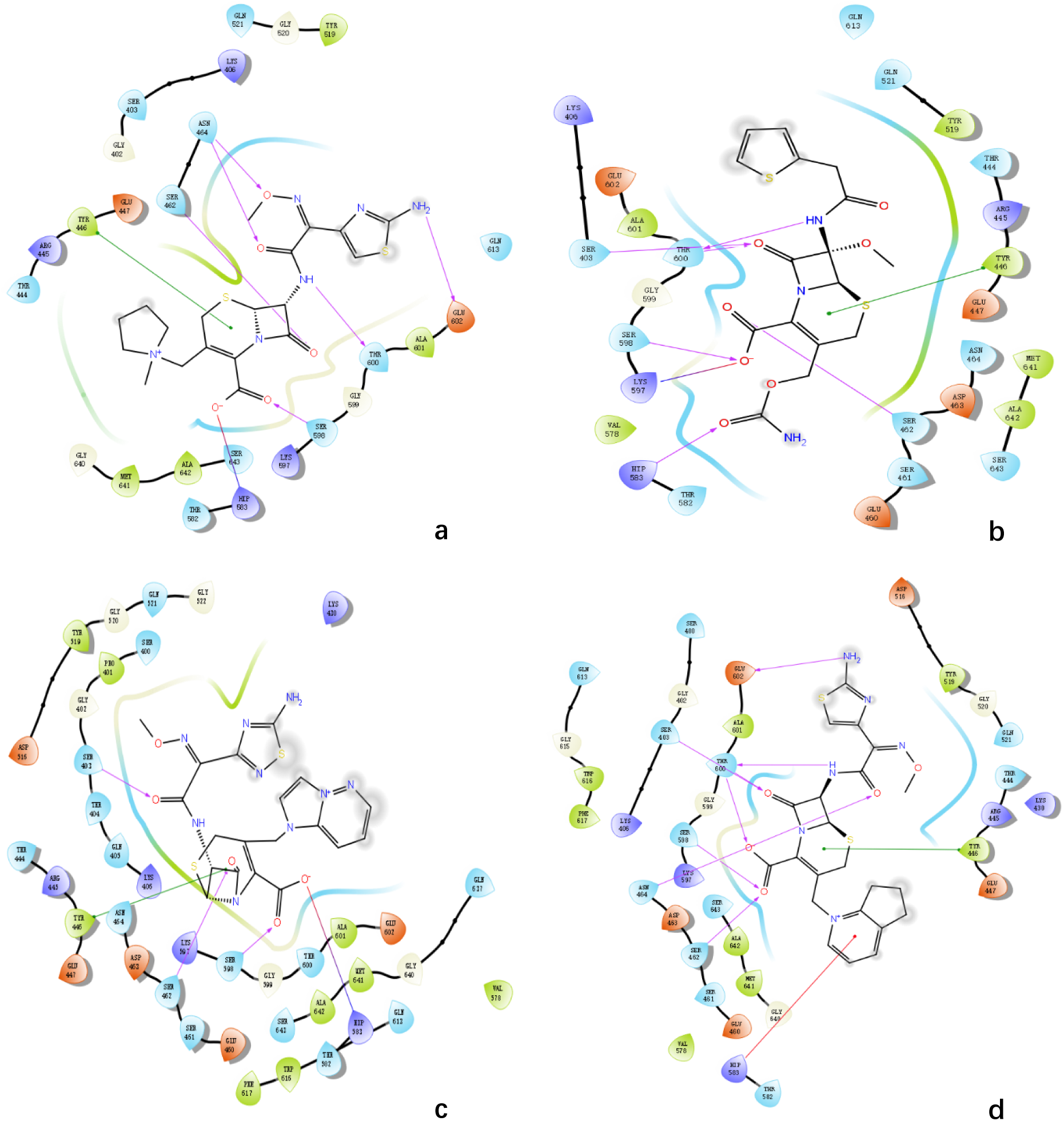
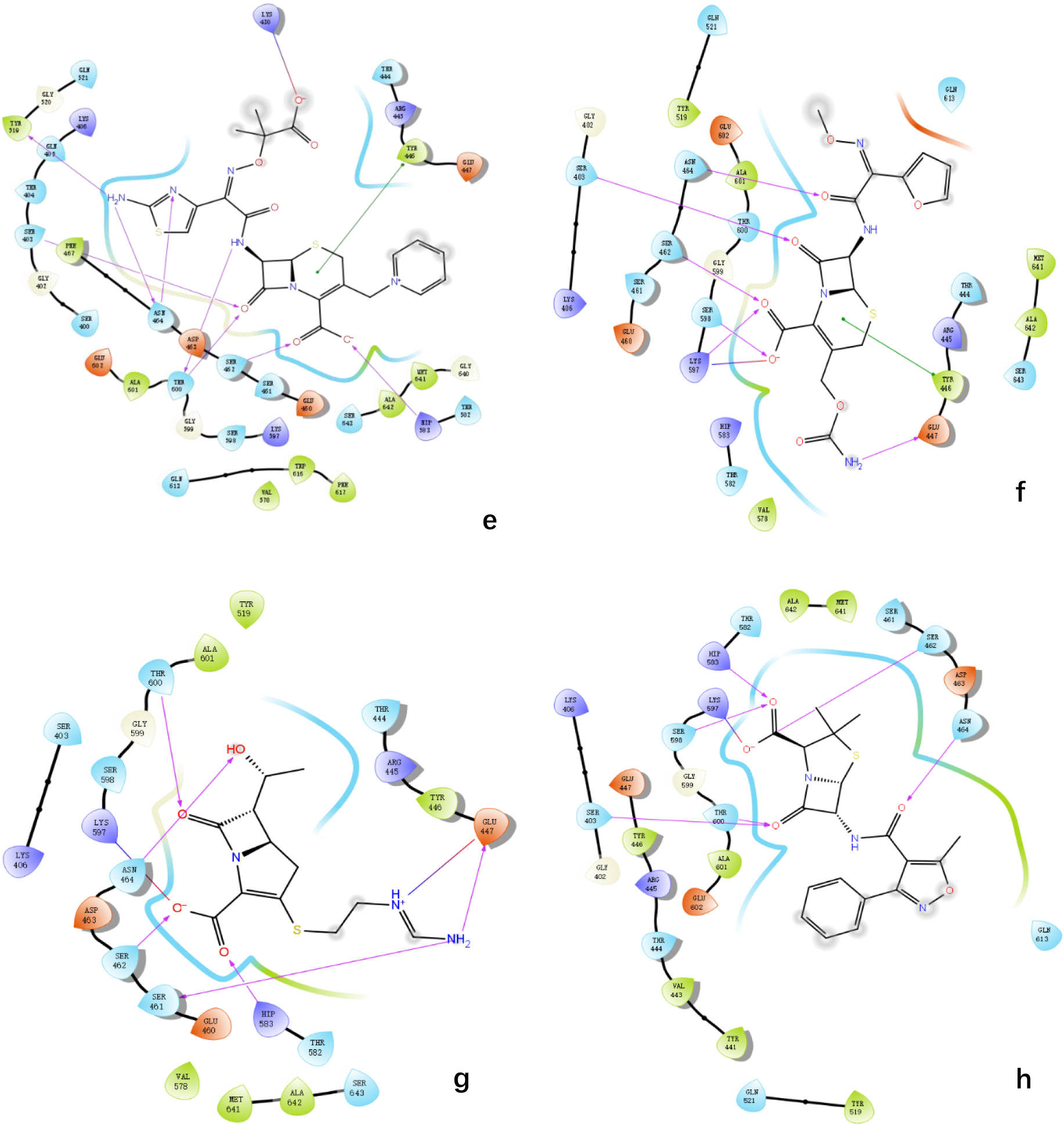
Analysis of non-covalent docking interaction patterns of different generations of β-lactam antibiotics. **a |**Interaction between 1MWU and Cefepime. **b |**Interaction between 1MWU and cefoxitin. **c |**Interaction between 1MWU and Cefozopran. **d |**Interaction between 1MWU and Cefpirome. **e |**Interaction between 1MWU and Ceftazidime. **f |**Interaction between 1MWU and Cefuroxime. **g |**Interaction between 1MWU and Imipenem. **h |**Interaction between 1MWU and oxacillin.

**Table 3.**
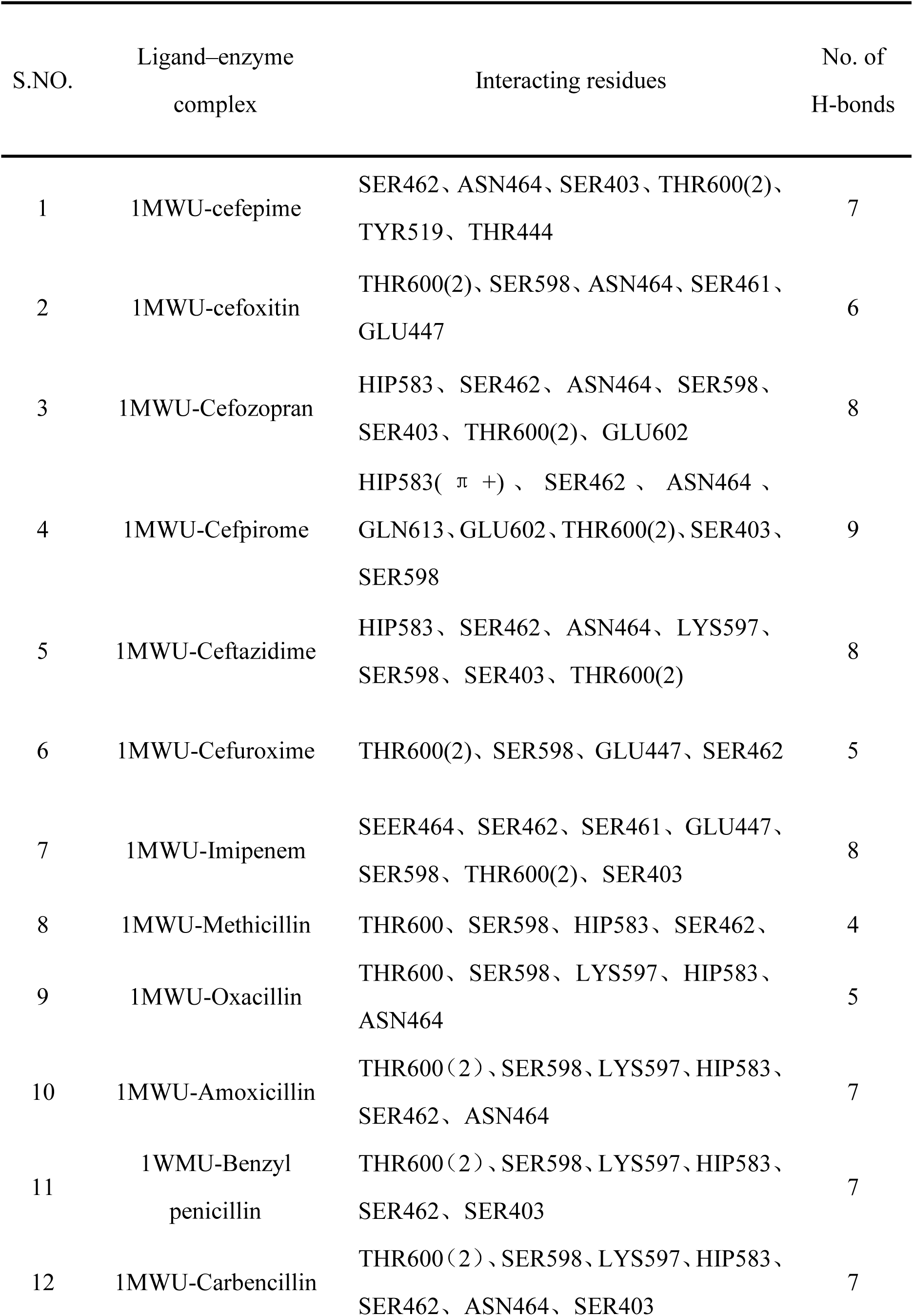
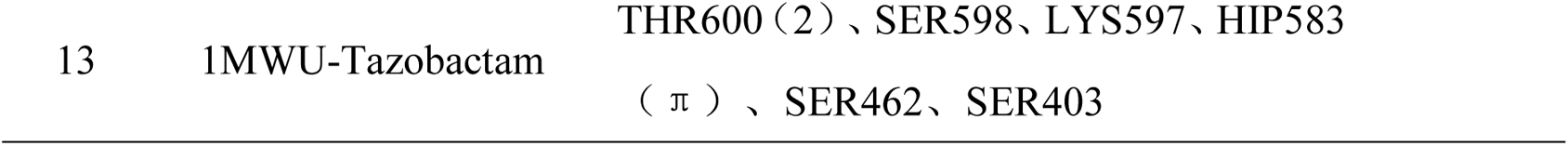
Covalent docking ofβ-lactam antibiotics with PBP2a.

**Table 4.**
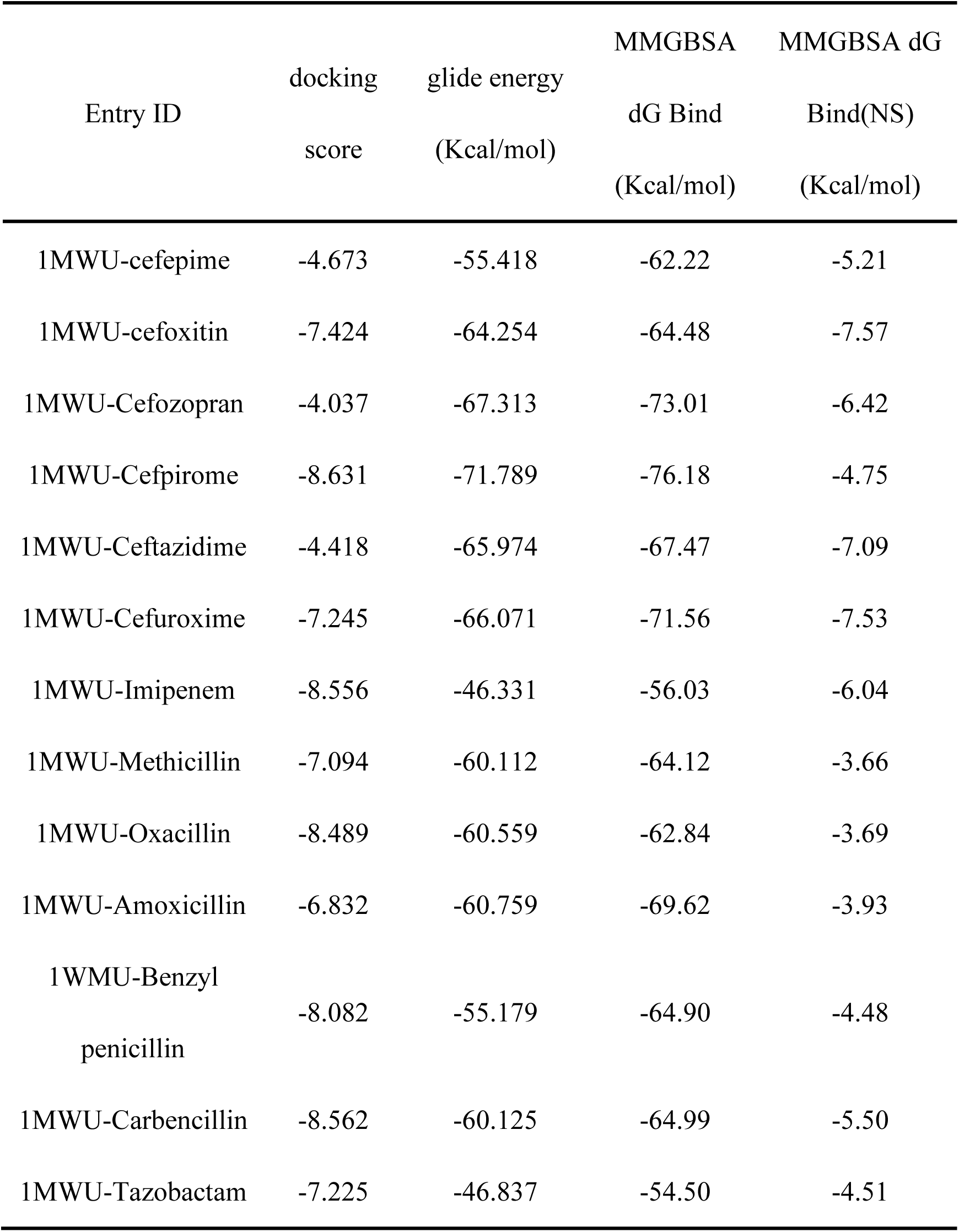
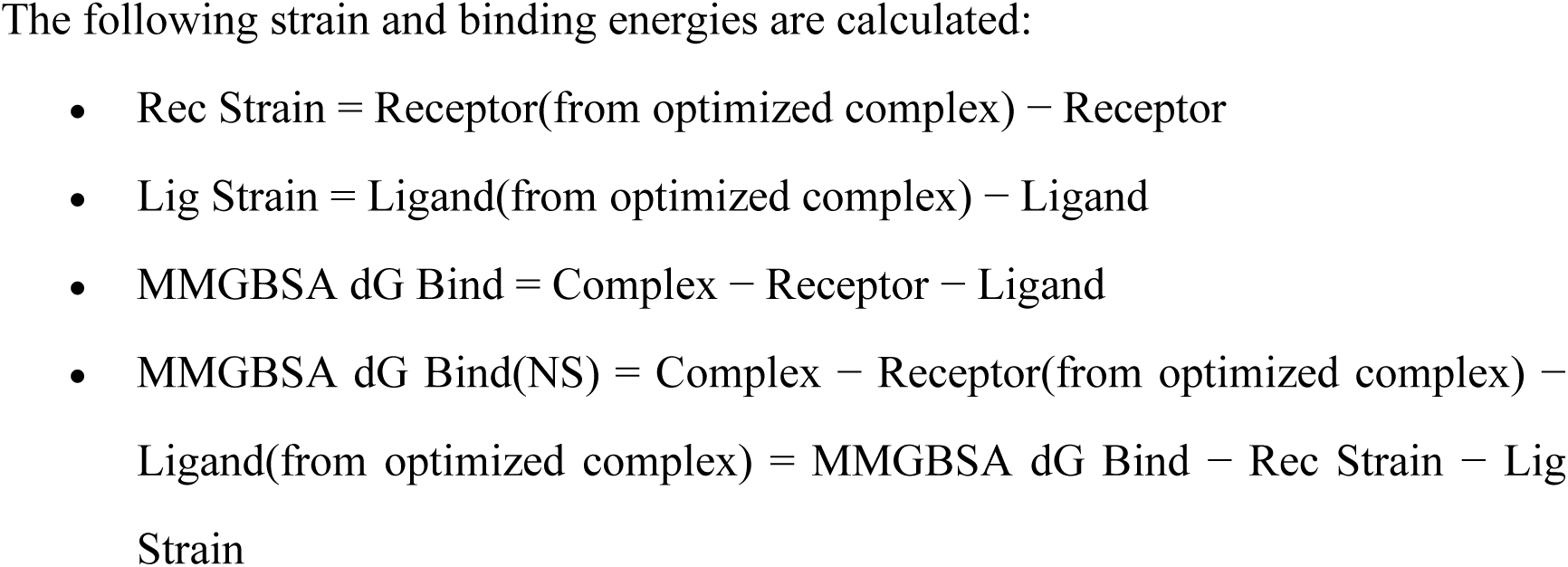
Analysis of covalent docking interactions between different generations of β-lactam antibiotics and PBP2a.

**Table 5.** Analysis of non-covalent docking interaction patterns of different generations of β-lactam antibiotics and PBP2a.

| S.NO. | Ligand–enzyme complex | Interacting residues | No. of H-bonds |
| --- | --- | --- | --- |
| 1 | 1MWU-cefepime | SER462、ASN462(2)、TYR446( $\pi$ )、HIP583、SER598、THR600、GLU602 | 8 |
| 2 | 1MWU-cefoxitin | THR600 (2) 、SER598、LYS597、HIP583、SER462、TYR446 ( $\pi$ ) 、SER403 | 8 |
| 3 | 1MWU-Cefozopran | SER403、TYR446 ( $\pi$ ) 、SER462、SER598、HIP583、 | 5 |
| 4 | 1MWU-Cefpirome | HIP583 ( $\pi$ ) 、SER462、ASN464、SER598、THR600 (3) 、GLU602、SER403、TYR446 | 10 |
| 5 | 1MWU-Ceftazidime | HIP583 、TYR446 、LYS430 、TYR519 、SER403、ASN464 (2) 、THR600 (2) 、SER462 | 10 |
| 6 | 1MWU-Cefuroxime | ASN464、SER403、SER462、SER598、LYS597 (2) 、GLU447、TYR446 | 8 |
| 7 | 1MWU-Imipenem | THR600 、LYS597 、HIP583 、ASN464 、SER462、SER461、GLU447 (2) | 8 |
| 8 | 1MWU-Methicillin | HIP583 、LYS597 、SER598 、THR600 、SER403、ASN464、SER462 | 7 |
| 9 | 1MWU-Oxacillin | THR600 、SER598 、LYS597 、HIP583 、ASN464、SER462、SER403 | 7 |
| 10 | 1MWU-Amoxicillin | THR600、SER598、LYS597、HIP583、SER462 | 5 |
| 11 | 1WMU-Benzyl penicillin | THR600、LYS597、HIP583、SER462、SER403 | 5 |
| 12 | 1MWU-Carbencillin | THR600 、SER598 、LYS597 、HIP583 、SER462、LYS430、SER403 | 7 |
| 13 | 1MWU-Tazobactam | LYS597、HIP583、SER462 | 3 |
| 14 | 1MWU-Meropenem | THR600 、SER598 、LYS597 、HIP583 、SER462、SER463、SER403 | 7 |

**Table 6.**
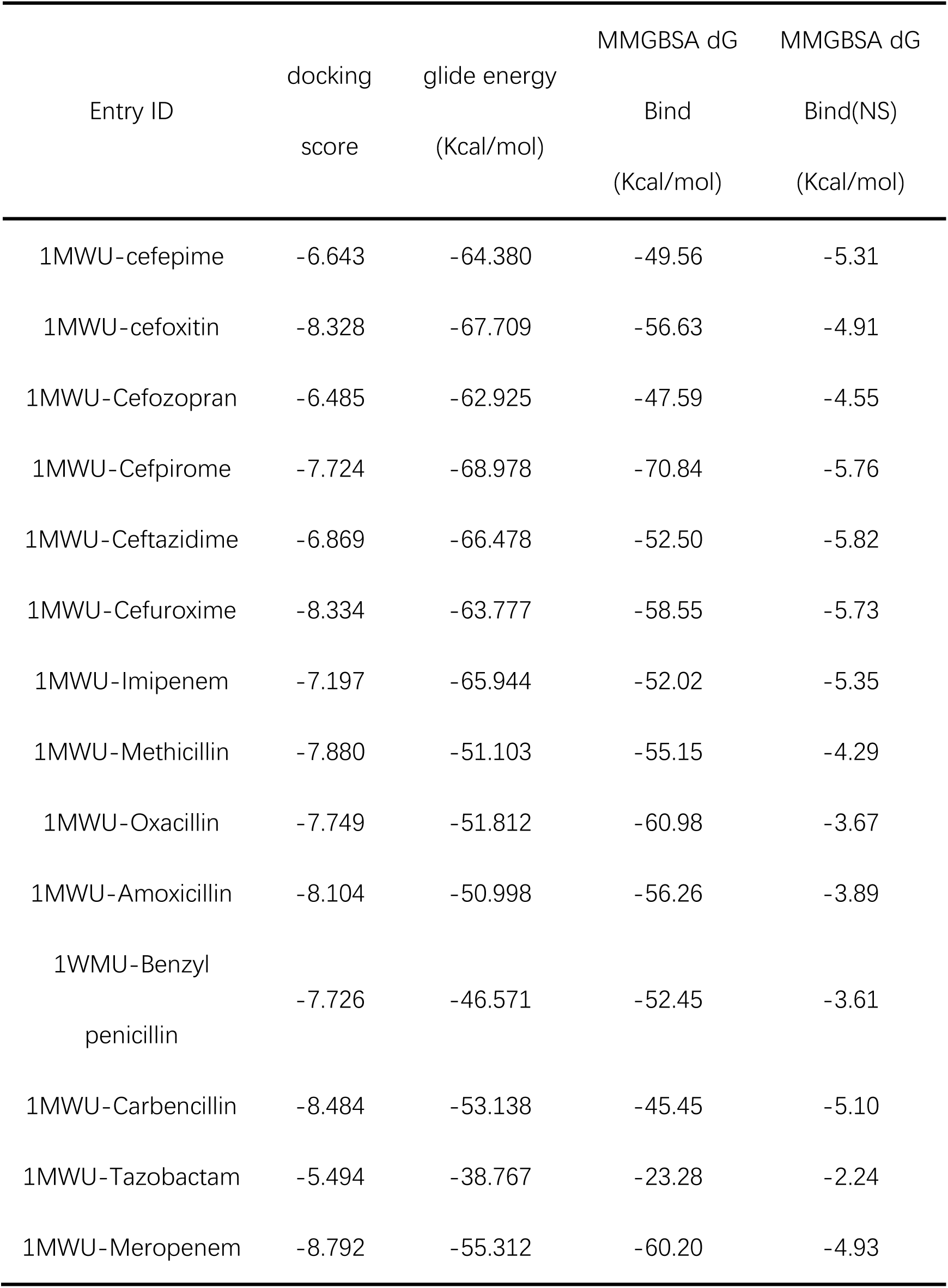
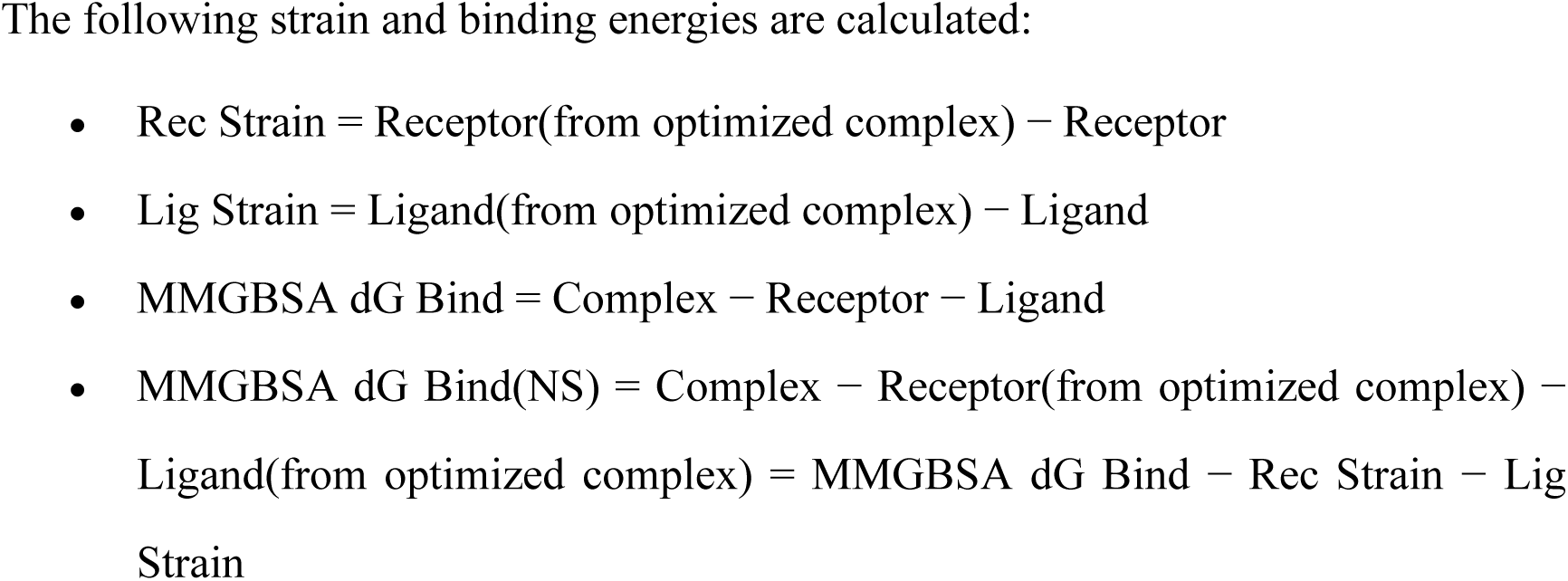
Non-covalent binding energy of different generations of β-lactam antibiotics to PBP2a.

**Table 7.** Molecular docking of β-lactam antibiotics with PBP2a.

| Ligand–enzyme<br>complex | Covalent docking |  |  | Non- covalent docking |  |  |
| --- | --- | --- | --- | --- | --- | --- |
|  | No. of<br>H-bonds | docking<br>score | MMGBSA<br>dG Bind<br>(Kcal/mol) | No. of<br>H-bonds | docking<br>score | MMGBSA<br>dG Bind<br>(Kcal/mol) |
| 1MWU-cefepime | 7 | -4.673 | -62.22 | 8 | -6.643 | -49.56 |
| 1MWU-cefoxitin | 6 | -7.424 | -64.48 | 8 | -8.328 | -56.63 |
| 1MWU-Cefozopran | 8 | -4.037 | -73.01 | 5 | -6.485 | -47.59 |
| 1MWU-Cefpirome | 9 | -8.631 | -76.18 | 10 | -7.724 | -70.84 |
| 1MWU-Ceftazidime | 8 | -4.418 | -67.47 | 10 | -6.869 | -52.50 |
| 1MWU-Cefuroxime | 5 | -7.245 | -71.56 | 8 | -8.334 | -58.55 |
| 1MWU-Imipenem | 8 | -8.556 | -56.03 | 8 | -7.197 | -52.02 |
| 1MWU-Methicillin | 4 | -7.094 | -64.12 | 7 | -7.880 | -55.15 |
| 1MWU-Oxacillin | 5 | -8.489 | -62.84 | 7 | -7.749 | -60.98 |
| 1MWU-Amoxicillin | 7 | -6.832 | -69.62 | 5 | -8.104 | -56.26 |
| 1WMU-Benzyl<br>penicillin | 7 | -8.082 | -64.90 | 5 | -7.726 | -52.45 |
| 1MWU-Carbencillin | 7 | -8.562 | -64.99 | 7 | -8.484 | -45.45 |
| 1MWU-Tazobactam | 6 | -8.102 | -71.43 | 3 | -5.494 | -23.28 |
| 1MWU-Meropenem | 6 | -7.968 | -70.99 | 7 | -8.792 | -60.20 |

- MMGBSA dG Bind = Complex – Receptor – Ligand

The drug that was showing higher binding energy and stronger interaction with protein was considered to be a drug with better susceptible activity^[39,41]^. The fourth and third generations of β-lactam antibiotics like cefpirome (-76.18 kcal/mol) and ceftazidime (-67.47 kcal/mol) were among the most effective antibiotic candidates against PBP2a. However, the drug that was showing lower binding energy and weaker interaction with protein was considered to be a drug with relative resistant capability. The resistance of PBP2a to the first and second generation of β-lactam antibiotics like benzyl penicillin (-52.45kcal/mol) and amoxicillin (-52.26kcal/mol) was also observed due to weaker molecular interactions and lower binding energies with the target.

Through systematic research, our findings indicate that there are quantifiable differences in the binding affinities of β-lactam antibiotics and PBP2a, that play a key regulatory role in drug resistance. Besides Ser403 as a covalent binding residue, Thr600, HIP583, Ser462, Asn464, and Ser598 were also identified as the essential amino acid residues, especially for cefpirome with a higher affinity to PBP2a. Thr600 was found to play an important role in binding to most of β-lactam antibiotics.

When dealing with the resistance of PBP2a, the main objective of β-lactam antibiotics is to target penicillin-binding protein 2 (PBP2) with precision. Covalent docking is particularly advantageous in cases where the specific binding site is identified. However, non-covalent docking is a computational method that enables ligands to autonomously navigate the whole surface of a target protein without prior knowledge of the directed site. Ligands have the ability to interact with many parts of the protein surface, rather than being limited to specific covalent binding sites. This enables a thorough examination of the whole protein surface near the active site. This aids in exploring alternate inhibiting areas outside established locations. This covalent docking research aligns with non-covalent docking to develop bioactive inhibitors that are both safe and efficient for targeting PBPs associated with MRSA.

### 3.3 Molecular dynamics simulation of PBP–β-lactams docking complex

#### 3.3.1 Root mean square deviation (RMSD) analysis of complexes

An in-depth investigation of molecular dynamics (MD) simulations using the Desmond package was conducted to examine the covalent docking results of PBP2a-β-lactam antibiotics complex. Tracking Root mean square deviation (RMSD) of proteins provides insight into their structural conformation throughout the simulation. RMSD represents the sum of all atomic deviations of the conformation from the target conformation at a given moment in time. RMSD analysis extracts valuable information about complex stability from ligand and protein molecule and can show whether the simulation reaches equilibrium-fluctuation at the end of the simulation.

The protein frames are initially aligned with the reference frame backbone, and then, the root-mean-square deviation (RMSD) is computed using the selected atoms. As shown in Figure 5, the RMSD of the complex system displayed progressive changes and leveled off after 60 ns. Proteins RMSD of PBP2a-b-lactam antibiotics complex may tolerate changes around the range of 2–5 Å without any issues. The simulation successfully reached convergence, displaying consistent RMSD values towards the end, indicating equilibration. Ligand RMSD of PBP2a-b-lactam antibiotics complex consistently stayed below 2.0 Å for a longer period of time,. This suggests that the ligand remains stable in the active site cavity of the protein.

**Figure 5.**
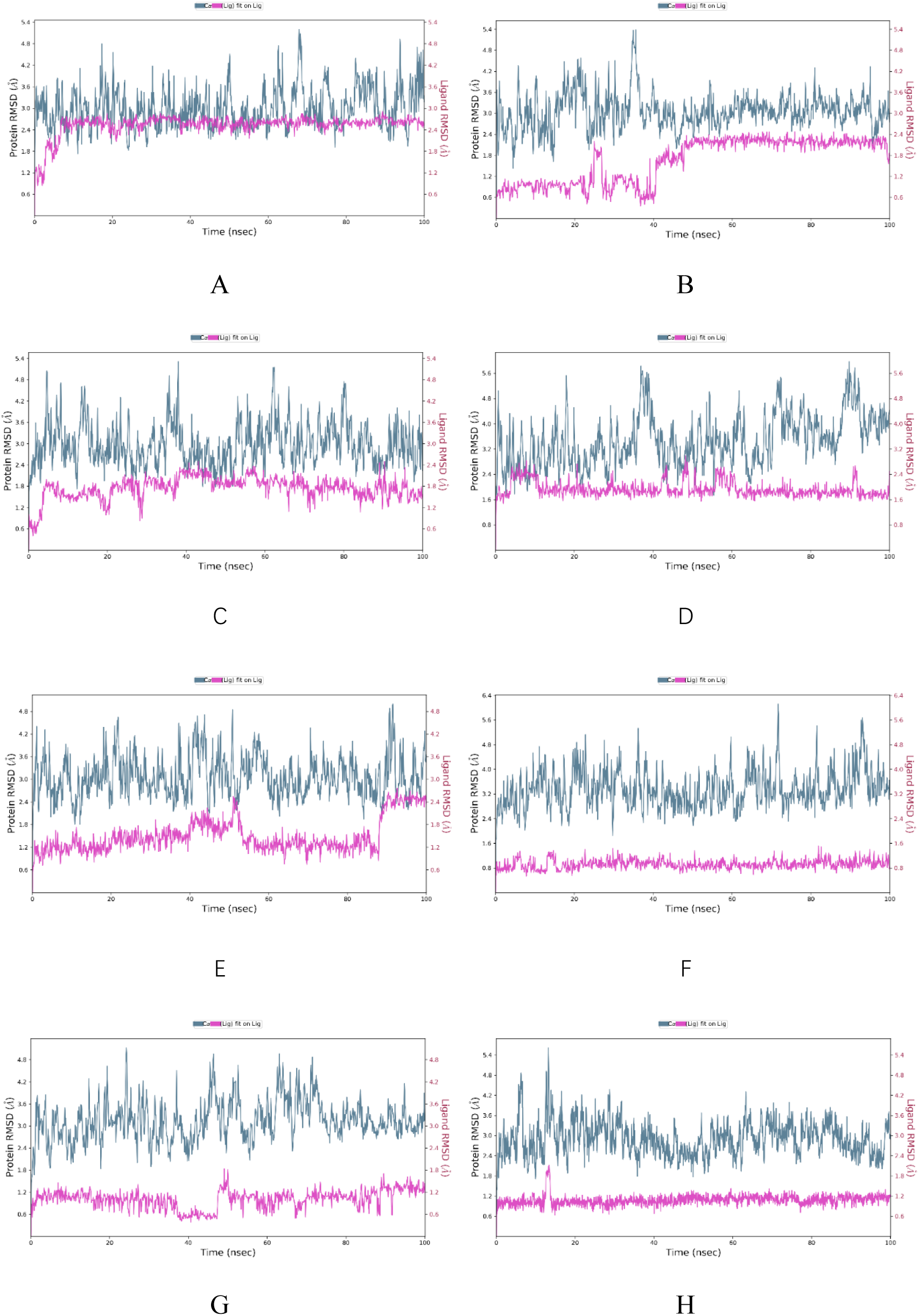
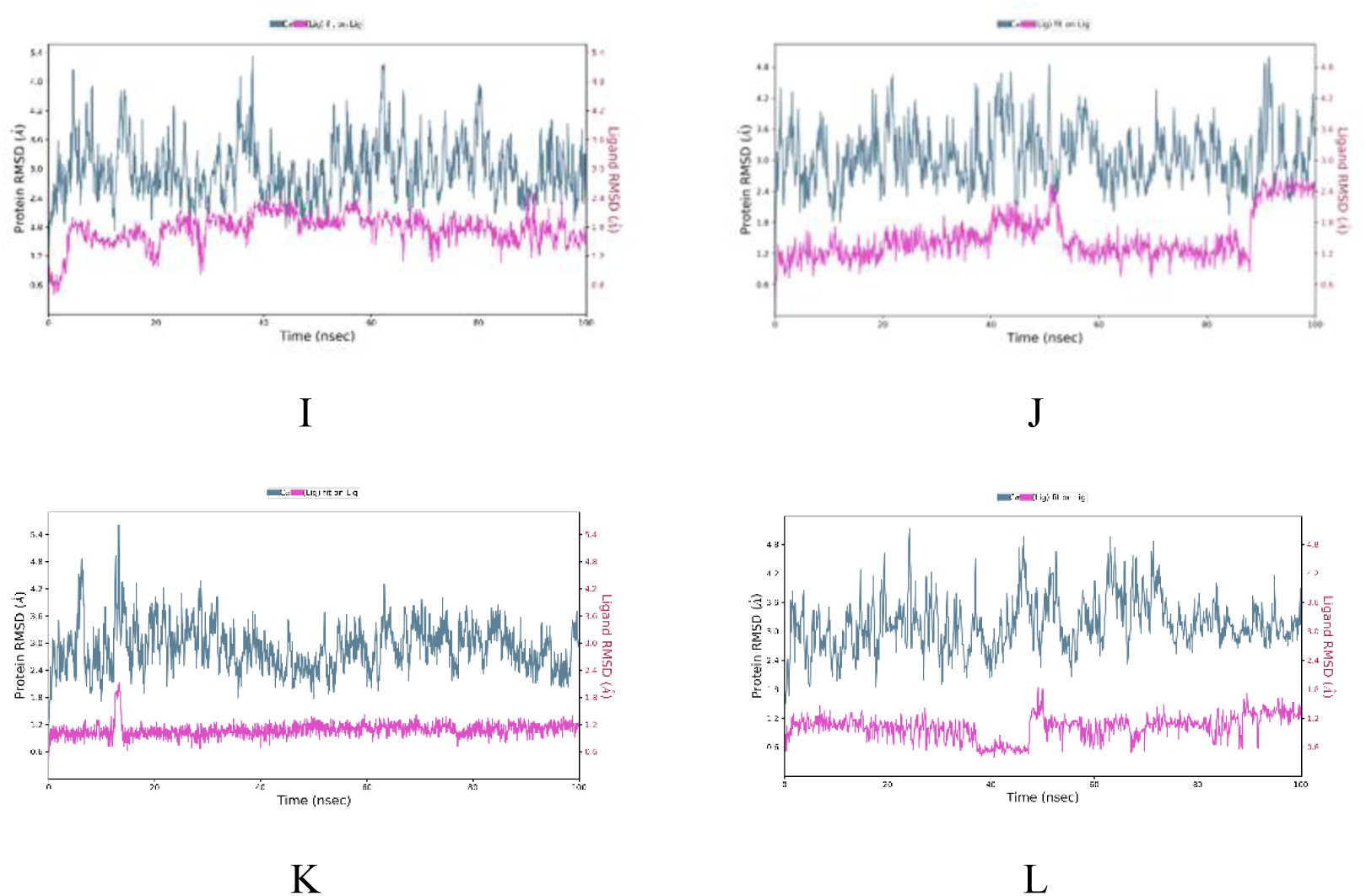
RMSD evolution of PBP2a-b-lactam antibiotics complex. The RMSD of proteins (left y-axis) considers only the C_α_ backbone atoms of the system, characterizing the displacement of the main chain atoms. The ligand RMSD (right y-axis) represents the degree of stabilization of the ligand connecting to the protein and its binding pocket, demonstrating the RMSD from the ligand backbone atoms, is able to give a clearer picture of the stability of the protein binding pocket. Blue color is the RMSD of the C_α_ backbone on the reference frame and red color is the RMSD of the ligand weight atoms. (A) Amoxicillin; (B) Benzylpenicillin; (C) Carbapenem; (D) Cefepime; (e) Cefoxitin; (F) Cefprozil; (G) Cefpirome; (H) Cefuroxime; (I) Cefozopran; (J) Ceftazidime; (K) Oxacillin; (L) Methicillin. Blue color indicates the protein (Cα main chain atom) of RMSD, and pink color indicates the RMSD of the lumenal ligand of the protein active site.

#### 3.3.2 Root Mean Square Fluctuation (RMSF) analysis

To further compare the effect of complex covalency on amino acid residues in the active site, Root mean square fluctuation (RMSF) analysis of the complexes was performed to quantify localized variations along the protein chain. RMSF can represent the flexibility of amino acids in proteins during simulations. If the RMSF of a certain region of the protein is significantly altered, it can be inferred that the ligand has affected that region of the protein, which may have induced a conformational change or stabilized the originally more flexible.

In Figure 6, RMSF from the trajectory file analyzes how the molecules attach to the active site of the protein, thus affecting the dynamics of the primary atoms. The more flexible amino acids have larger RMSF values, lower flexibility amino acids have lower RMSF values and peaks indicate protein regions that fluctuate the most. Typically, the tails (N- and C-termini) are observed to fluctuate more than the rest of the protein. Secondary structural elements Structural elements (e.g., α-helices and β-strands) are usually relatively more stable than the unstructured portion of the protein and therefore fluctuate less than the cyclic region. The ligand contacts section offers significant insights into the protein residues that interact with the ligand. Understanding the particular residues involved in ligand interactions enhances our overall comprehension of the dynamics of protein-ligand binding during the simulation. According to the calculations, most of the amino acids have RMSF values below 2, indicating that the overall flexibility of the complex is low.

**Figure 6.**
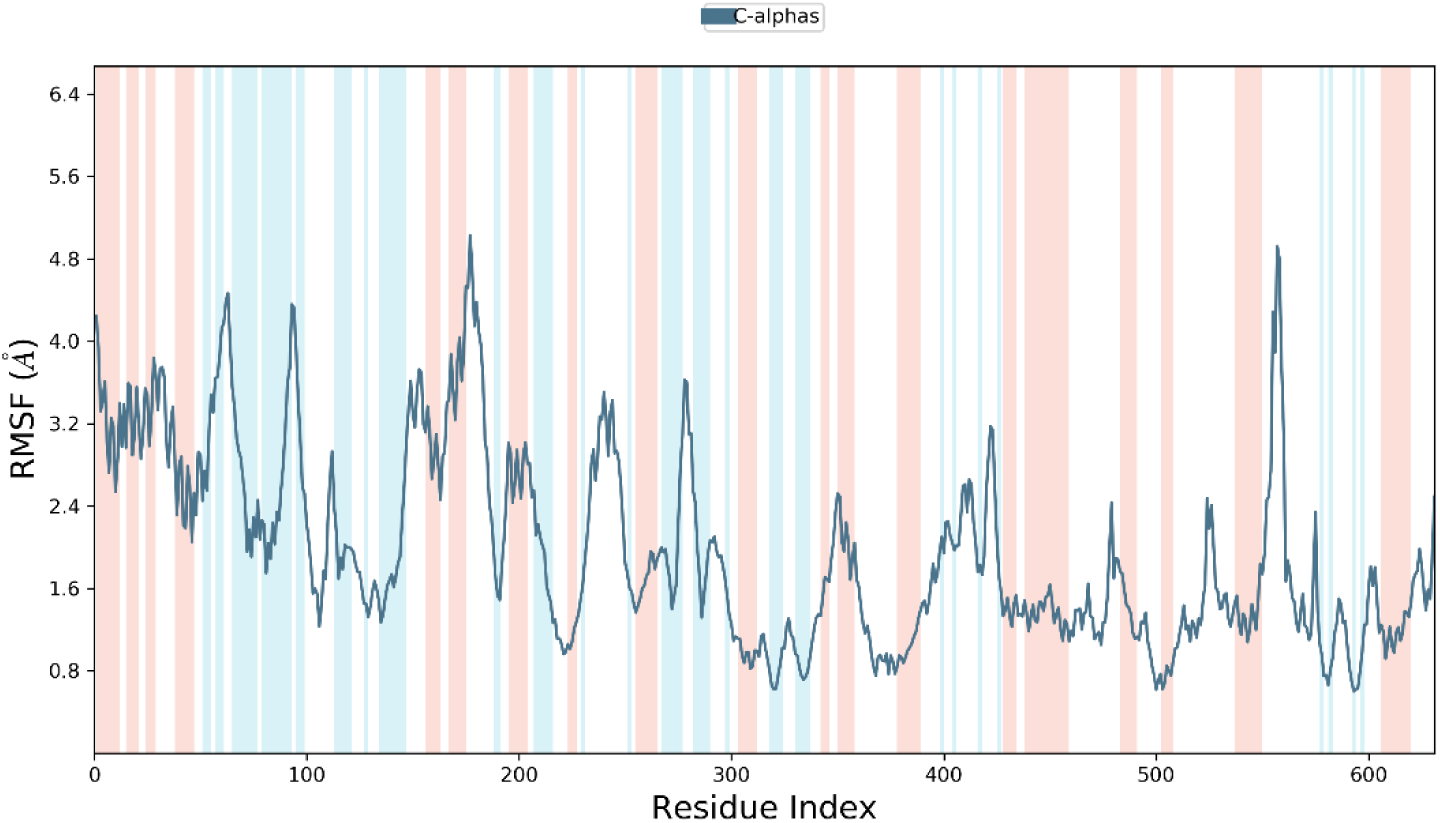
Graph Root Mean Square Fluctuation (RMSF) can be used to characterize local variations in protein chains (left y-axis). X-axis is the position of residues, and the blue and red stripes in the graph are the secondary structure of proteins, with the blue representing β-folding and the red representing α-helices, which characterize the local RMSF variations in protein chains (left y-axis).

#### 3.5.3 Hydrogen bond analysis of the protein-ligand complex

Because Hydrogen bonding plays a crucial role in the stability of protein-ligand complexes and also has a great impact on drug specificity, MD simulation was employed to investigate the strength of hydrogen bonding for different ligands. Figure 7 provides the timeline representation of interactions and contacts (H bonds) across the simulation trajectory. The Proteins formed many hydrogen bonds with the ligands throughout 100 ns MD simulation, suggesting they were relatively tightly bound.

**Figure 7.**
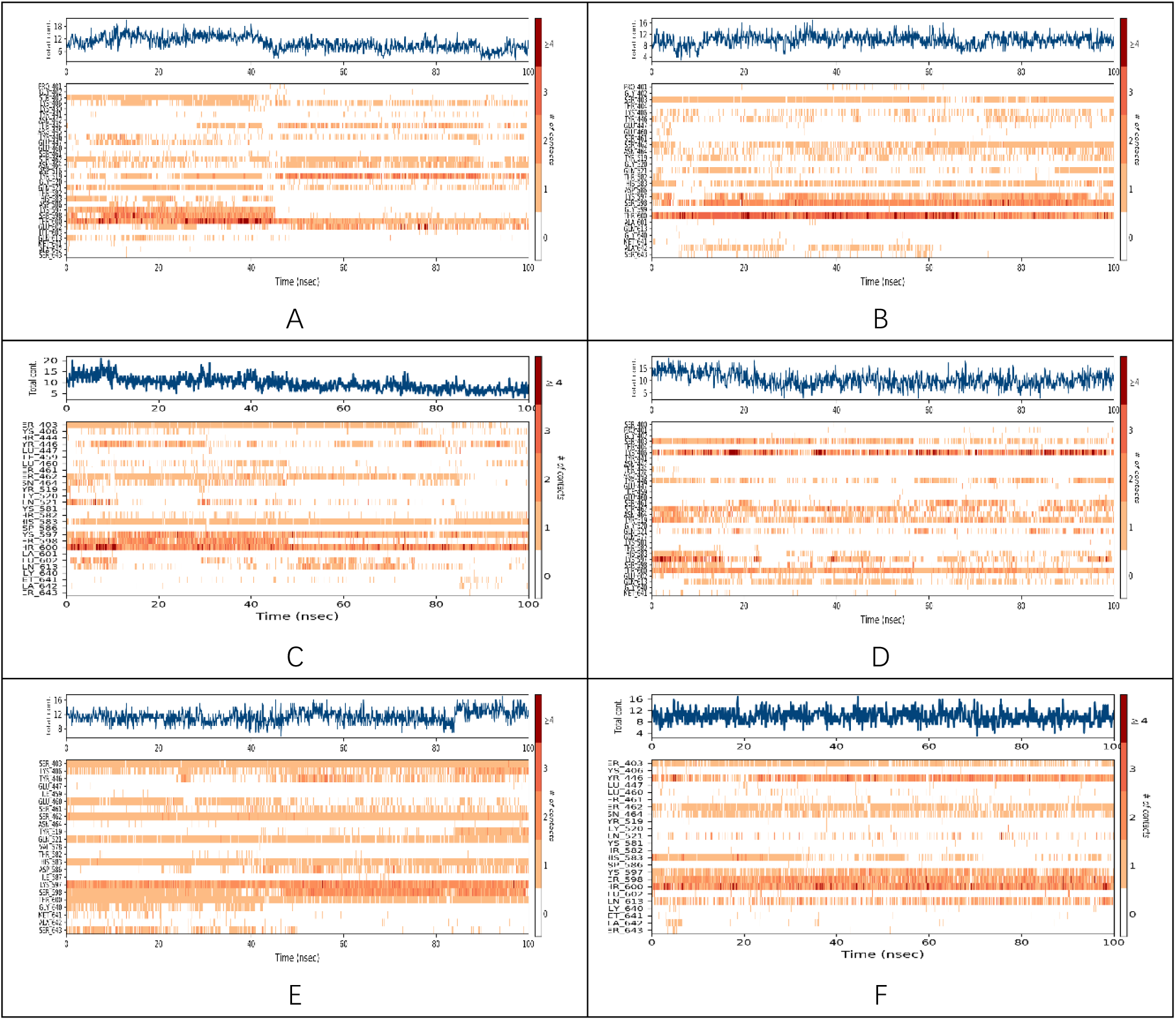
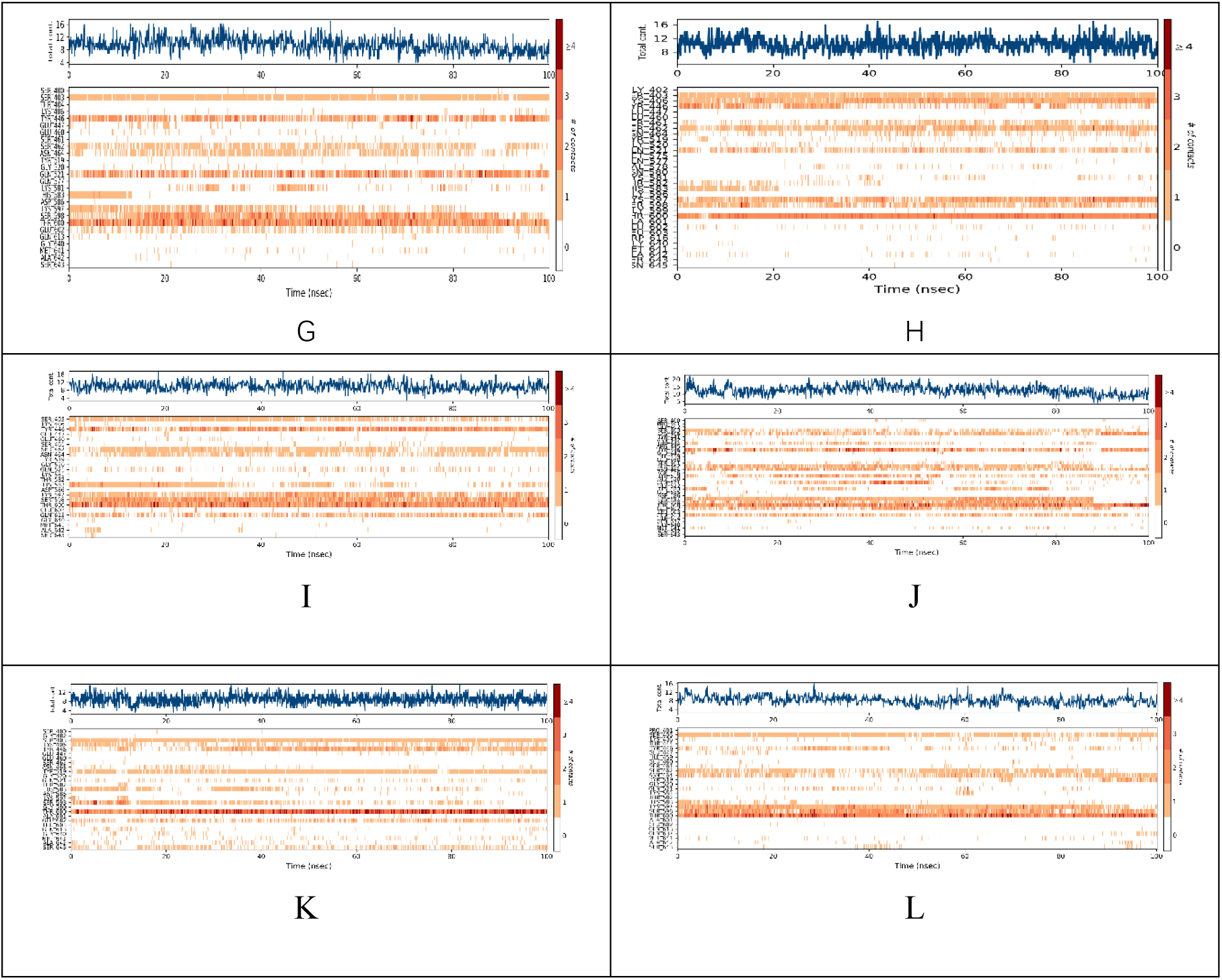
Timeline representation of interactions and contacts (H bonds). (A) Amoxicillin. (B) benzy penicillin. (C) cabencillin. (D) cefepime. (E) cefexitiin. (F) cefopran. (G) cefpirome. (H) cefuroxime. (I) Cefozopran; (J) Ceftazidime; (K) Oxacillin; (L) Methicillin.

The top panel shows the total number of specific contacts that the protein makes with the ligand throughout the entire trajectory. The graph shows the residues that interact with the ligand in each trajectory frame interacting with the ligand. Some residues have more than one specific contact with the ligand, and these are indicated by dark orange shading, with darker orange shading indicating more contacts, according to the scale on the right side of the graph.

The timeline representation provided a concise overview of the specific contacts and revealed sustained contacts. These findings collectively give a clear picture of the stable binding interface. THR600, SER598, HIP583 and SER462 all showed prominent hydrogen bonding interactions, which were maintained until the end of the simulation. Cefexitin and Cefpirome had stronger hydrogen bonding interactions with PBP2a, which suggests that these two antibiotics will probably play a better role in dealing with PBP2a.

## 4 Conclusion

Deep understanding of the intricate molecular mechanisms involved in antibiotic resistance is of utmost importance. This study examines the influence of a series of β-lactam antibiotics on the molecular interaction with PBP2a using both covalent and non-covalent docking methods based on prior knowledge of penicillin-resistant and penicillin-susceptible. These specific antibiotics have been discovered to interfere with the active site, resulting in challenges for susceptible or resistant changes during β-lactam antibiotic acylation.

Advanced analytical techniques were employed to simulate the binding process and evaluate interactions. Molecular docking analyses investigate essential residues, highlighting exceptional binding affinity and their potential for designing an inhibitor of PBP2a. The binding affinity of four generations of b-lactam antibiotics is highlighted through molecular docking analyses, suggesting a promising candidate of PBP2a inhibitor. Molecular dynamics simulations offer a detailed comprehension of the interactions between antibiotics and PBP2a. The verified simulation outcomes establish a basis for the creation of potent inhibitors and medicinal therapies to combat infectious illnesses

